# The spatial distribution and temporal trends of livestock damages caused by wolves in Europe

**DOI:** 10.1101/2022.07.12.499715

**Authors:** Liam Singer, Xenia Wietlisbach, Raffael Hickisch, Eva Maria Schoell, Christoph Leuenberger, Angela Van den Broek, Manon Désalme, Koen Driesen, Mari Lyly, Francesca Marucco, Miroslav Kutal, Nives Pagon, Cristian Remus Papp, Paraskevi Milioni, Remigijus Uzdras, Ilgvars Zihmanis, Fridolin Zimmermann, Katrina Marsden, Klaus Hackländer, José Vicente López-Bao, Sybille Klenzendorf, Daniel Wegmann

**Affiliations:** Department of Biology, Université de Fribourg, Chemin du Musée 15, CH-1700 Fribourg, Switzerland; Swiss Institute of Bioinformatics, CH-1700 Fribourg, Switzerland; Department of Mathematics, Université de Fribourg, Chemin du Musée 20, CH-1700 Fribourg, Switzerland; EuroLargeCarnivores: https://wiwraj.eurolargecarnivores.eu/en/; University of Natural Resources and Life Sciences, Vienna (BOKU), Department of Integrative Biology and Biodiversity Research, Institute of Wildlife Biology and Game Management, Gregor-Mendel-Strasse 33, 1180 Vienna, Austria; BIJ12, Leidseveer 2, 3511 SB Utrecht, the Netherlands; Direction Régionale de l’Environnement, de l’Aménagement et du Logement d’Auvergne-Rhône-Alpes, France; Agency for nature and forests of the Flemish Government, Belgium; Finnish Wildlife Agency, Kampusranta 9C, FI-60320, Seinäjoki, Finland; University of Torino, Department of Life Sciences and Systems Biology - DBIOS, Via Accademia Albertina 13, 10123 Torino, Italy; Department of Forest Ecology, Faculty of Forestry and Wood Technology, Mendel University in Brno, 61300 Brno, Czech Republic; Friends of the Earth Czech Republic, Carnivore Conservation Programme, Dolní náměstí 38, 779 00 Olomouc, Czech Republic; Slovenia Forest Service, Večna pot 2, SI-1000 Ljubljana, Slovenia; Department of Taxonomy and Ecology, Faculty of Biology and Geology, Babeş-Bolyai University, 5-7 Clinicilor Street, 400006 Cluj-Napoca, Romania; Hellenic Agricultural Insurances Organization, Greek Ministry of Agriculture and Food, Greece; State Service for Protected Areas under the Ministry of Environment, Antakalnio g. 25, LT-10312 Vilnius, Lithuania; State Forest Service of Latvia, 13. janvāra iela 15, LV-1932, Latvia; Stiftung KORA, Talgut-Zentrum 5, 3063 Ittigen, Switzerland; Adelphi consult GmbH, Alt-Moabit 91, 10559 Berlin; Biodiversity Research Institute (CSIC - Oviedo University - Principality of Asturias), Oviedo University, E-33600 Mieres, Spain; WWF Germany, Reinhardtstr. 18, 10117 Berlin; Deutsche Wildtier Stiftung (German Wildlife Foundation), Christoph-Probst-Weg 4, 20251 Hamburg, Germany

**Keywords:** wolf, livestock predation, human–wildlife conflict, damage trends, Europe

## Abstract

Wolf populations are recovering and expanding across Europe, causing conflicts with livestock owners. To mitigate these conflicts and reduce livestock damages, authorities spend considerable resources to compensate damages, support damage prevention measures, and manage wolf populations. However, the effectiveness of these measures remains largely unknown, especially at larger geographic scales. Here we compiled incident-based livestock damage data across 21 countries for the years 2018, 2019 and 2020, during which 39,262 wolf-caused incidents were reported from 470 administrative regions. We found substantial regional variation in all aspects of the data, including the primary target species, the density of damages, their seasonal distribution, and their temporal trend. More than half of the variation in damage densities across regions is explained by the area of extensively cultivated habitats occupied by wolves and namely natural grasslands and broad-leaved forests. Regional variation in husbandry practices and damage prevention, while difficult to quantify at a continental scale, appear important factors to further modulate these incidents. As illustrated with detailed data from Germany, for instance, the relationship between the number of wolf units and damages is diminishing over time, suggesting some adaptation of livestock owners and local authorities to their presence, for example by increasing prevention efforts. As we argue, temporal trends of damage incidents, which are robust to variation in data collection across regions, are thus informative about the local intensity of the wolf-human conflict. We estimated increasing trends for the majority of regions, reflecting the current expansion of wolves across the continent. Nonetheless, many of these increases were moderate and for more than one third of all regions, trends were negative despite growing wolf populations, thus indicating that wolf-livestock conflicts can be successfully mitigated with proper management.

## 1. Introduction

The last decades have seen the recovery of wolves (*Canis lupus*) across Europe, including in several regions where the species had previously been extinct for decades or even centuries (Chapron et al., 2014; Reinhardt et al., 2019). Between 2012 and 2016, an estimated 17,000 wolves roamed the European continent (excluding Russia and Belarus, Boitani et al. (2018)) and, with the exception of one isolated population in Spain (López-Bao et al., 2018), all populations are continuing to expand (Chapron et al., 2014; Linnell and Cretois, 2018). This recolonization process is taking place without reintroductions and is due to three main factors: first, wolves are granted strict legal protection in many countries by the EU Habitats Directive and/or the Bern Convention (Chapron et al., 2014; Epstein et al., 2016). Second, populations of important prey species such as roe deer, red deer and wild boar were able to recover following land abandonment and reforestation in Europe (Trouwborst, 2010). Third, wolves have a remarkably high adaptive capacity, allowing them to establish in fragmented, human-dominated landscapes (Mech and Boitani, 2007; Trouwborst, 2010; Sazatornil et al., 2016; Cimatti et al., 2021).

Wolf recovery is not exempt from social tensions and conflicts (Dressel et al., 2015; Skogen et al., 2017). While ecologists are regarding the growing wolf population in Europe as a conservation success story, many farmers in recovering areas fear increased depredation of their livestock and, as a consequence, a threat to their livelihoods (van Eeden et al., 2018; Bautista et al., 2019; Rode et al., 2021). It is imperative to address these conflicts and to facilitate the coexistence of humans and wolves to ensure positive conservation outcomes in a Europe densely populated by humans. The absence of wolves for an extended period has often resulted in reduced adaptations for coexistence (López-Bao et al., 2017), which in turn harbors potential for conflict once the species is recolonizing its former habitat (Chapron et al., 2014; Gervasi et al., 2021a).

To mitigate these conflicts, authorities aim at raising the standards of livestock protection to reduce livestock vulnerability and shift depredation from livestock to wild prey (van Eeden et al., 2018; Eklund et al., 2017). In Europe, most countries provide financial support to their farmers to procure and maintain livestock damage prevention measures such as electric fences, livestock guarding dogs, or permanent herding, either via the Common Agricultural Policy (CAP) (Marsden and Hovardas, 2020) or similar schemes (e.g. Agridea, 2022). Lethal interventions, both illegal and by governments to remove problem individuals (Ordiz et al., 2013), are additional elements of damage prevention, yet their efficiency remain controversial due to a lack of empirical, conclusive evidence (Santiago-Avila et al., 2020; Bruns et al., 2020). Along with the varying quality and scope of implementation of non-lethal measures has given way to a debate on whether and what kind of damage prevention performs best (Eklund et al., 2017; Bonnet et al., 2019; Oliveira et al., 2021). A large-scale randomized control trial on the effectiveness of different prevention measures is still missing to date (van Eeden et al., 2018).

To shed more light on our understanding of livestock damages caused by wolves in Europe, several studies attempted to identify factors explaining their spatial variation, yet often with conflicting results. At regional scales, for instance, several studies reported that livestock in heterogeneous landscapes and in particular close to forest edges were the most vulnerable to wolf predation (Rigg et al., 2011; Kaartinen et al., 2009b). Across multiple countries, however, no landscape features were found to correlate with the number of compensated sheep (Gervasi et al., 2021a). In contrast, the number of wolves correlated positively with the number of compensated sheep at the scale of multiple countries (Gervasi et al., 2021a), yet at regional scales, incidents were reported to increase with the geographic spread of wolves, but not with an increase in their numbers (Khorozyan and Heurich, 2022).

In an attempt to reconcile these findings, we compiled a large European-wide data set of incident-based livestock damage incidents at the municipality level from 2018-2020. We then characterized their distribution in space and time and examined the extent to which regional densities in damages can be explained by wolf presence and landscape features reflecting the density and overlap between wolves and livestock. We further estimated regional trends in damage incidents, which we argue are helpful indicators for coexistence of wolves and humans.

## 2. Material and Methods

### 2.1. Case-based livestock damage incidents

We collected case-based livestock damage data for 2018, 2019 and 2020 at the regional level (i.e. NUTS3 regions, see below), where cases are incidents of livestock depredation as recorded by authorities. While most reported incidents reflect a single attack of wolves on livestock, they may rarely involve multiple attacks if livestock was not checked daily. To obtain case-based data, we consulted the websites of regional authorities if available. Otherwise, we reached out to regional and national authorities of all EU member states, Norway and Switzerland (Supplementary Table S.1) in spring 2019, 2020 and 2021 to report livestock damage incidents of the previous year using a template questionnaire (Supplementary Table S.2). Contacts were mediated by a national collaborator from the EU Life EuroLargeCarnivores programme. Our questionnaire consisted of fixed-response questions to be filled per incident, with an option to comment in a separate column. The main attributes were (i) the primary asset missing, injured or killed, (ii) the assessment level or probability of the cause being identified correctly, (iii) the amount of compensation paid per incident, and (iv) the damage prevention measure implemented at the time of the incident in the broad categories defined by Eklund et al. (2017): *electric fence, wire fence, livestock guarding dog, permanent shepherd*, and *other qualified protection*.

We translated the submitted information to English. If the questionnaire was returned incomplete, we followed up and entered any additional information we received by hand. While many respondents adhered to our fixed-answer request, some replies had to be curated manually to match our standards: (i) If more than one asset species was reported for the same incident, we recorded the incident for each species separately, but kept the same incident ID and treated the event as a single incident in our analyses; (ii) if no assessment level on the certainty of wolf predation was reported, we recontacted the authorities for clarification. In case we did not receive any information, we chose the category *unspecified*. (iii) If more than one date was reported for the same incident, we took the first reported date. (iv) If no geographic coordinates were submitted indicating the location of the damage incident, we used the village name (or the smallest geographic unit available) and converted it into geographic coordinates using the Google Geocoding API (Google, 2022). If neither geographic coordinates nor geographic units were given, or if the provided name could not be converted to coordinates, we removed the incident from our analysis.

As we accumulated data annually, we sent along a report including descriptive statistics as well as the finalized national data set of the previous year for cross-checking by the authorities. In addition, we shared initial exploration of the data to demonstrate the relevance of such data.

### 2.2. Geographical regions

We conducted our analyses at different geographic scales: at the continental and country level, as well as for the three levels of the Nomenclature of Territorial Units for Statistics (NUTS; European Commission and Eurostat, 2020) that subdivide each country into smaller geographic units. The NUTS regions mostly follow the administrative subdivisions of the EU Member States (see Figure 1 for a visualization). In addition, they are unambiguously standardized across Europe and are strictly hierarchical: each country is composed of one or more NUTS1 region, each of which are composed of one or more NUTS2 regions and so forth. Benefiting from this hierarchical setup, we first compiled the counts n_ik_ of reported damage incidents for each year *Y_k_* ∈ {2018, 2019, 2020} and all 470 NUTS3 regions *i* = 1,…, *I*. We next obtained counts *n_rk_* = ∑_*i*∈*r*_ *n_ik_* for all geographic regions *r* in NUTS1, NUTS2, each country and the continent as a whole by summing across all NUTS3 regions *i* encompassed in *r*, denoted here as *i* ∈ *r*.

**Figure 1:**
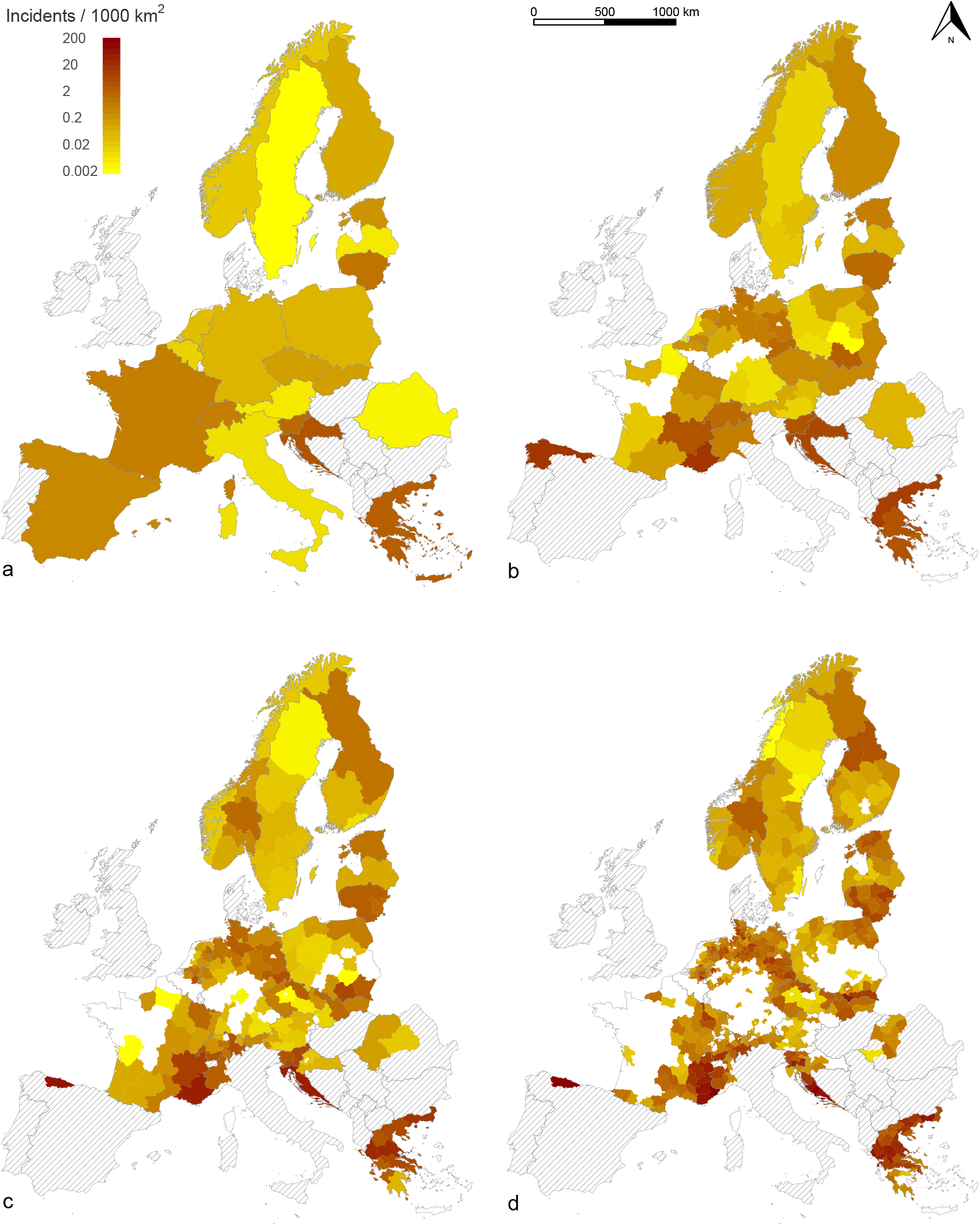
Average wolf-caused livestock incident density plotted on the country level (a), NUTS1 level (b), NUTS2 level (c) and NUTS3 level (d) across the years 2018, 2019 and 2020. Gray shaded regions indicate regions from which we did not obtain data. Regions shown in white indicate regions from which no damages were reported.

We restricted these counts to incidents for which wolves were sufficiently likely the cause: for administrative regions that provided an assessment level, we kept those with category *presumed correct* or *confirmed*. For administrative regions that did not provide an assessment level, we considered only incidents for which a compensation was paid. If neither was provided, we considered all submitted incidents assuming that only sufficiently probable incidents were shared with us.

### 2.3. Seasonal distribution of damage incidents

To characterize the seasonal distribution of damage incidents, we aggregated all available incidents by month, discarding all incidents for which no date was provided. To test for temporal variation, we performed *χ*^2^-tests on monthly counts versus their expectation under a uniform distribution. To test if incidents generally occur later in the year in northern than southern Europe, we performed Mann-Whitney-U tests on the months, grouping all incidents reported from Estonia, Finland, Latvia, Lithuania, Norway and Sweden (north) and Croatia, France, Greece, Italy and Spain (south). All tests were performed with the statistical software R (R Core Team, 2021), using the functions `chisq.test()` and `wilcox.test()`.

### 2.4. Covariates explaining wolf-caused damage incidents

We investigated whether covariates may explain the variation in damage incidents between NUTS3 regions. To account for the non-independence of annual damage incidents across years, we focused on average annual counts *n_r_*. We used the average rather than the sum to account for the few NUTS3 regions for which we obtained data for two years only.

#### 2.4.1. Considered covariates

We considered the following ten covariates.

- **Area of wolf presence** To characterize wolf presence across Europe, we used the shapefile compiled for the most recent period available (2012 to 2016) at a 10 x 10 km resolution for the Large Carnivore Initiative of Europe IUCN Specialist Group and for the IUCN Red List Assessment (Kaczensky et al., 2021). This map encompasses the entire region considered in this study. At each grid point, the authors translated the presence and frequency of wolves into one of three categorical variables: *permanent*, *sporadic* or *no presence*. We intersected this map with the NUTS3 regions using the *st-intersection()* function from the *sf* package (Pebesma, 2018) in R (R Core Team, 2021) and used this intersection to determine the area (in km^2^) permanently and sporadically occupied by wolves for each NUTS3 region, denoted by *Per-A* and *Spo-A*, respectively.
- **Wolf area by land cover classes** We used the CORINE Land Cover (CLC, code 18) data from the year 2018 (European Environment Agency, 2018) to quantify land cover for all analyzed regions. The available data covers our area of interest at a scale of 1:100,000. The classification comprises artificial surfaces, agricultural areas, forests and semi-natural areas, wetlands and water bodies. We used the `st_intersection()` function from the `sf` package (Pebesma, 2018) in the statistical software `R` (R Core Team, 2021) to intersect the CLC layers with the area occupied by wolves (either permanently or sporadically, see above), and further with each NUTS3 region. This way we obtained for each NUTS3 region and as a proxy for suitable wolf refuge areas the area occupied by wolves of broad-leaved (*BLF-A*) coniferous (C-A) and mixed forests (*MF-A*), along with the area occupied by wolves of pastures (*P-A*) and natural grasslands *NG-A* aas a proxy for wolf hunting areas as well as livestock presence and availability.
- **Historical continuity of wolf presence** Following Gervasi et al. (2021a) and based on previous estimates (Chapron et al., 2014), we determined for each NUTS3 region, whether (1) or not (0) wolves were present during the 1950-1970s, a factor denoted by *50ya*.
- **Grazing season length** The mean per regions of the bioclimatic variable *BIO11* of WorldClim (Fick and Hijmans, 2017), which was previously found to be a good predictor of the grazing season length across Europe, explaining 52% of the total variation (Phelan et al., 2016).
- **Support for prevention measures** To quantify policies regarding the support of livestock damage prevention schemes, we used an ordinal variable *Prev* stating whether prevention measures were financially supported between 2018-2020 in a given political region (yes, *partially*, or *no*) as defined in Marsden and Hovardas (2020). Our data relates to support for the purchase of fencing and livestock guarding dogs, as data on their funding is most commonly available. A partial support means that the financial support provided does not cover the full costs of the prevention measures. We used information from Marsden and Hovardas (2020) for Croatia, Finland, France, Greece, Latvia and Lithuania. For the remaining regions we used information provided by the European commission (European Union, 2022), or additional publications.

To test whether we missed any major environmental factor, we also explored models that considered for each region i) the latitude of the centroid, ii) the average altitude as provided by the Digital Elevation Model for Europe (dowloaded from https://www.mapsforeurope.org/datasets/euro-dem on November 2, 2022) and iii) the mean of each bioclimatic variable available from WorldClim (BIO1 through BIO19) after transforming all temperatures to Kelvin.

#### 2.4.2. Considered models

Let us denote by *z_rc_, c* = 1,…,*C* and *f_rd_, d* = 1,…, *D* sets of numerical and factor covariates for each region *r*, respectively. We considered two types of models to account for the heteroscedasticity present in the data: A Poisson model with log-link function of the form

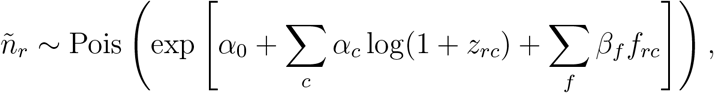

where 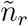 denotes the average annual counts *n_r_* rounded to the nearest integer, *α*_0_ an intercept and *α_c_* and *β_f_* the regression coefficients. All numerical covariates (e.g. area occupied by wolves) were log-transformed to maintain their expected linear relationship with incidents counts, but we added one to each to allow zero values.

To avoid the need of rounding and log-transforming covariates, we also fitted a Gaussian model with power-transform of the form

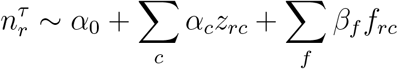

where *τ* denotes the parameters of the power transform. We chose the value of *τ* that maximized the variance explained of the full model using all covariates, and identified it using a line search.

To identify the best sub-models (i.e. selection of covariates), we used the function `dredge()` from the package muMin (Barton and Barton, 2015) in R, used the Akaike Information Criterion (AIC) to identify the best models, and used the function `anova()` from the package *stats* (Bates et al., 1992) to determine the fraction of the total variation explained by each model.

### 2.5. Trends in livestock damages

We estimated trends in livestock damage incidents for each region *r* using a Bayesian inference approach similar to that in Aebischer et al. (2020), but extended to more than two time points. Let *n_ik_* denote the observed incident counts in NUTS3 region *i* in year *Y_k_*, *k* = 1,…, *K*. We assumed these counts are Poisson distributed *n_ik_ ~* Poisson (*λ_ik_*s_i_) with means proportional to two region-specific factors: the rate *λ_ik_* at which incidents occur in the region *i* during year *Y_k_*, and rate *s_i_* with which incidents are reported in that region.

We assumed that for any region *r*, incident rates follow a common exponential trend with rate *γ_r_* such that *λ_ik_* = *λ*_*i*0_ exp(*Y_k_γ_r_*) for all *i* ∈ *r*, and we sought to infer the rate of change *γ_r_* from all incident counts *n_ik_* reported for all NUTS3 regions *i* ∈ *r* for all years *k*. To do so, we conditioned on the total numbers of counts *ν_i_* = ∑_*k*_ *n_ik_* across years for each NUTS3 region (see Link and Sauer, 1997). The conditional distribution of ***n**_i_* = (*n*_*i*1_,…, *n*_*iK*_) given *ν_i_* is multinomial:

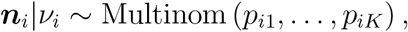

(Johnson et al., 1997) in our case with probabilities

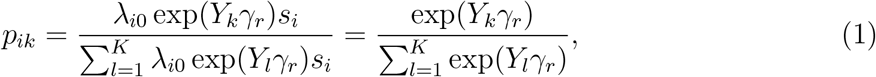

where the sums runs across all years *l* = 1,…,*K*. Due to conditioning, the nuisance parameters *λ*_*i*0_ and *s_i_* are canceled out from the fraction, rendering trend estimates independent of any variation in reporting rates across administrative regions.

The likelihood of the full observation vector ***n*** = (***n**_i_*, *i* ∈ *r*), conditional on ***ν*** = (*ν_i_*, *i* ∈ *r*), is

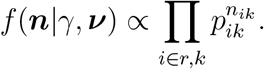

Following Aebischer et al. (2020), we chose the non-informative Jeffrey’s prior for *γ_r_*, which is (up to a normalizing constant) the square root of the determinant of the Fisher information

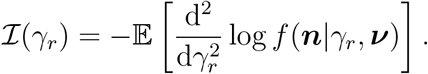

Using 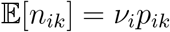 and ∑_*k*_*P_ik_* = 1, we arrive at

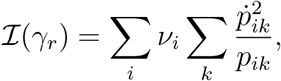

where

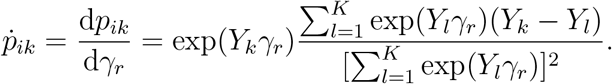

We implemented an Markov chain Monte Carlo (MCMC) approach to generate samples from the posterior 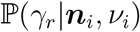 under this model in an R package (see data availability statement). Using the function `birp_data()`, we created one data set per NUTS region for which damage incidents were reported for at least two years (setting all efforts to 1.0) and then inferred trends for this region using the function `birp()`.

We then classified each region as having an increasing or decreasing trend if the posterior mode 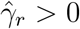 or 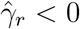 respectively, and quantified the uncertainty *p_γ__r_* associated with these point estimates as the posterior probability indicating the opposite sign:

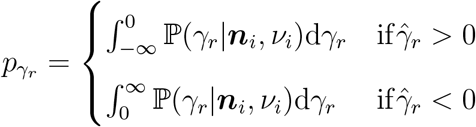

Using the above method, we inferred trends for the total number of incidents combined across all affected species as well as for each species individually. Pearson correlations among the species-specific trend estimates were calculated using the function `cor.test in R` (R Core Team, 2021), restricting the calculation to regions for which trends could be estimated for both species.

To test if trends where higher in regions only recently colonized, we used a Mann-Whitney U test on the posterior modes against either *50ya* or a factor indicating whether a NUTS3 region was bordering other regions without damage incidents (1) or not (0).

### 2.6. Testing for spatial autocorrelation

We used Moran’s *I* to test for autocorrelation in the estimated damage trends (posterior modes) and other metrics across each NUTS3 region. For each pair of regions *i* and *j*, we used a weight *w_ij_* = 0,1 indicating whether the regions share a common border (*w_ij_* = 1) or not (*w_ij_* = 0), assessed using the function `st_touches()` from the package `sf` (Pebesma, 2018) in R (R Core Team, 2021). We assessed the significance of *I* against a null distribution obtained by permuting the values randomly across regions one million times.

To test for variation on a north-south cline, we further determined the latitude of the centroid of each NUTS3 region using the function `st_centroid` of the `sf` package.

### 2.7. Explaining variation in livestock damage incident trends between regions

For Germany, information on wolf occurrences is available at a finer spatial and temporal scale from (BIJ12 et al., 2022) since 2000. We mapped these occurrences to NUTS regions with `st_intersection` as above and calculated the number of known wolf units *w_rk_* per NUTS3 region *r* for the years *Y_k_* ∈ {2018, 2019, 2020}. In contrast to previous analyses (e.g. Reinhardt et al., 2019), we treated wolf individuals, pairs and packs each as one territorial unit since single individuals may also cause extensive damage.

We then used the function `cor.test()` in `R` to test for correlations between the number of wolf units in each region *w_rk_* and the number of reported incidents *n_rk_*, limiting the analyses to regions for which wolves, incidents or both were reported. We further used these data to test for correlations between trends in livestock damage incidents and trends in wolf occurrences. For this, we inferred trends in the number of known wolf units 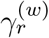 for each NUTS region *r* using the same approach as described above for incidents. We then tested for correlations between the trends inferred for wolf units 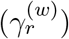 and those inferred for damage incidents (*γ_r_*) at all NUTS levels, only considering regions for which at least one wolf was reported in any of the years 2018, 2019 or 2020. Finally, we tested if the inferred trends correlate with the number of years that wolves were present in each region, defined as the number of years between the first reported wolf and 2020.

### 2.8. Data availability

The data that supports the finding of this study, namely the full records of damage incidents as well as all environmental covariates used to model them, are openly available at Zenodo (the double-blind review process of Biological Conservation does not allow us to share the DOI prior to acceptance). Note that some information considered sensitive by the authorities was withheld (compensation payments and/or exact geolocation data), but can be obtained by contacting authorities directly (Supplementary Table S.1). The above Zenodo link further includes all R code written to conduct our analyses, with the exception of the R package to perform the trend analyses, which is available at (the double-blind review process of Biological Conservation does not allow us to share the DOI prior to acceptance).

## 3. Results

We collected data on livestock damage incidents caused by wolves for 2018, 2019 and 2020 from national or regional authorities for the following 16 countries: Austria, Belgium, Croatia, Czech Republic, Finland, France, Germany, Greece, Latvia, Netherlands, Norway, Poland, Slovakia, Slovenia, Sweden and Switzerland. We obtained partial data for five additional countries: For Estonia we could only obtain data for 2018 and 2019, and for Lithuania only for 2019 and 2020. For Italy, Romania and Spain, we received data only for a subset of the provinces (seven, eight and one NUTS3 regions, respectively). Requests were declined or left unanswered by five countries: Belarus, Denmark, Hungary, Portugal and Ukraine. In total, we obtained data for 910 NUTS3 regions, of which 470 reported incidents. The total number of reported incidents was 43,703, of which 43,513 (99.6%) could be unambiguously attributed to a single NUTS3 region and were kept for our analyses. These incidents were distributed as 13,895, 15,086 and 14,532 across the three years 2018, 2019, 2020, respectively.

We further restricted our analyses to incidents for which wolves were sufficiently likely the cause. A subset of countries (Austria, Belgium, Switzerland, Germany, Estonia, Greece, Croatia, Latvia, Netherlands, Norway, Poland, Sweden, Slovenia) provided an assessment level. Of the 22,112 incidents from these countries, we kept 17,904 (80.97%) that were reported as *confirmed* (4,803, 21.7%) or *presumed correct* (13,101, 59.2%), and excluded incidents that were *negative* (2,017, 9.1%), *uncertain* (706, 3.2%), *no assessment possible* (1,200, 5.4%), *assessment pending* (269, 1.2%) or *unspecified* (16, 0.1%). For Finland, Romania and Spain that did not provide an assessment level, we kept the 9,181 (99.5%) incidents for which compensation was paid. For the remaining countries (Czech Republic, France, Italy, Lithuania, Slovakia) that provided neither information, we kept all 12,177 incidents, assuming that only sufficiently probable incidents were shared with us. In total, we thus kept 39,262 (89.8%) incidents (Supplementary Table S.3) and will refer to these as *wolf-caused incidents* below.

The countries with the highest number of reported wolf-caused incidents across the three years were France (9,840), Greece (6,870) and Spain (6,856). The countries with the lowest number of reported wolf-caused incidents were Belgium (79), Latvia (91) and Austria (115). As shown in Figure 1, regions varied greatly in their densities of wolf-caused incidents, with south-eastern France, coastal Croatia, northern Greece and the Spanish province of Asturias being regional hotspots of livestock damage incidents in our data set.

Most data collected was not associated with the information on the application of damage prevention measures (84.3%), with only eight countries (Belgium, Croatia, Germany, Latvia, Netherlands, Poland, Slovenia, Sweden) reporting whether or not a prevention measure was applied at the time of the incident. Among those incidents (6,158, 15.7%), the most common measure was *electric fence* (767 incidents, 12%), followed by *wire fence* (311, 5%), *guarding dog* (31, 0.5%) and *permanent shepherd* (8, 0.13%). For an additional 3,826 (62%) of incidents the prevention measure was indicated as *other qualified protection*, while 1,430 (23%) affected unprotected animals. Note that for 224 incidents (3.6%), multiple measures were in place.

### 3.1. The species most frequently targeted by wolves

In terms of wolf-caused incidents, sheep were most frequently affected (21,301, 54.2%), followed by cattle (7,672, 19.5%) and goats (4,328, 11%). Other animals less frequently affected included horses (3,125 wolf-caused incidents, 8%), reindeer (1,976, 5%), dogs (529, 1.4%), domestic deer (red, roe or fallow deer, 201, 0.5%), donkeys (166, 0.4%), pigs (10, <0.1%) and lamas or alpaca (8, <0.1%). For 343 (0.9%) additional incidents, the affected animals were not indicated to the species level. For Finland, the most affected species was reindeer (85.8%), for Greece cattle (46.5%) and for Spain (Asturias) horses (42.3%). For the remaining 18 countries, the most frequently affected species was sheep (46.0-97.6%).

Across the 39,262 wolf-caused incidents reported from all 21 countries, 99,056 animals were killed, injured or went missing. The distribution of the number of affected animals per incident was heavily skewed with 58.9% of all incidents involving a single animal, 23.2% involving two or three and only 3.5% involving ten or more individuals. Only two incidents involved more than 100 animals, the largest being the only reported incident from the Romanian province of Timis affecting 402 animals. Sheep had the most causalities (71,023, 71.7%), and goats had more casualties (11,338, 11.4%) than cattle (8,415, 8.5%) in line with goat incidents usually involving more animals (2.6 on average, standard deviation 2.6) than cattle incidents (1.1 on average, standard deviation 0.6). Incidents involving sheep involved 3.3 animals on average (standard deviation 5.2).

### 3.2. Seasonal distribution of wolf-caused incidents

To gain insights into seasonal patterns of livestock damage incidences, we aggregated records by months, discarding 359 (0.9%) incidents which did not provide a date. As shown in Figure 3, incidents show strong temporal variation (*χ*^2^ = 4480.9, p < 10^-15^). Across all species, incidents peak between July and October with 48.7% of the total incidents falling within these months. This pattern is particularly visible for sheep (55.2%), as well as for cattle (43.4%) and goats (40.9%), albeit less pronounced. In contrast, incidents involving horses peak between April and July (51.8%) and those involving reindeer between September and December (67.5%).

**Figure 2:**
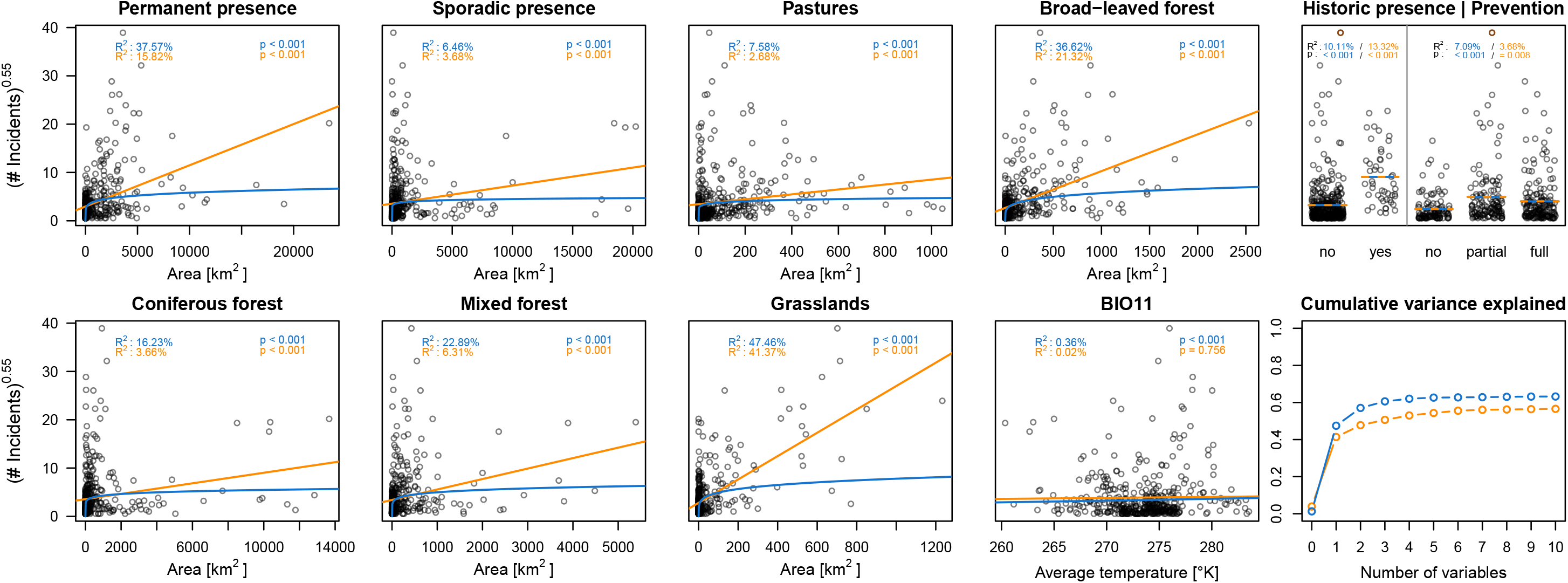
Correlation of ten environmental covariates and the number of reported wolf-caused livestock damage incidents in our data set, excluding the outlier region of Asturias (ES120). The Poisson model is shown in blue, the Gaussian model is shown in orange. For visualization purposes, both models are plotted using the same power-transformation on the incidents.

**Figure 3:**
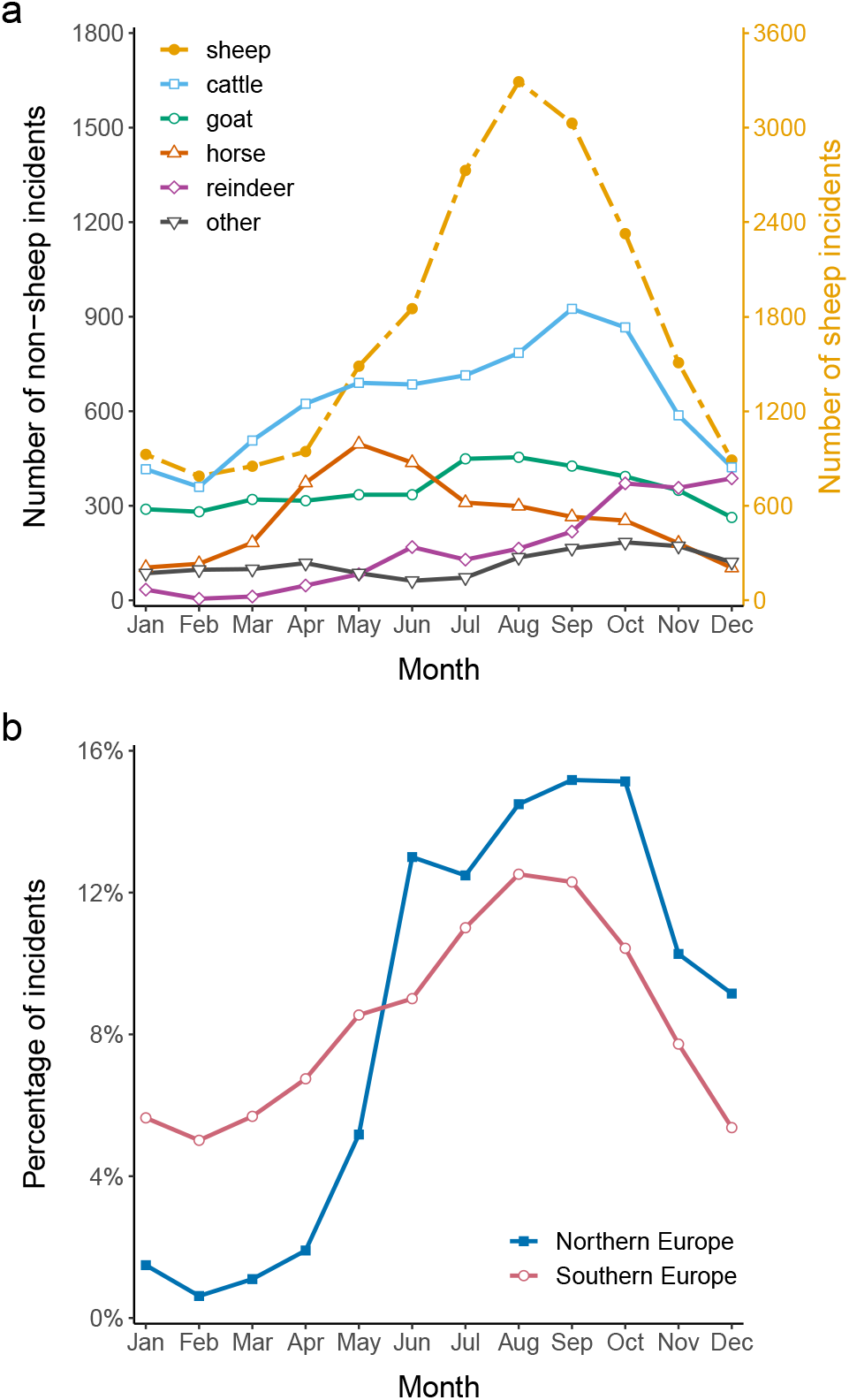
(a) Monthly distribution of wolf-caused livestock damage incidents. (b) Percentage of monthly wolf-caused livestock damage incidents for northern Europe (Estonia, Finland, Latvia, Lithuania, Norway, Sweden) and southern Europe (Croatia, France, Greece, Italy, Spain).

The differences between target species are explained by the geographic distribution of livestock and the observation that incidents in northern Europe (Estonia, Finland, Latvia, Lithuania, Norway and Sweden) generally occur later in the year than in southern Europe (Croatia, France, Greece, Italy and Spain, *U* = 51011510, p < 2.210^-16^): in southern Europe, 23.3% of all incidents occur before May, but in northern Europe only 5.1%. This pattern was also found for each species with enough data to perform the Mann-Whitney-U test, namely sheep (*U* = 14902386, p < 0.042), cattle (*U* = 678569, p < 10^-8^), goats (*U* = 87644, p < 10^-3^), horses (*U* = 1410.5, p < 10^-3^) and dogs (*U* = 22569, p < 10^-4^).

### 3.3. Covariates explaining wolf-caused damage incidents

The livestock species affected by wolf depredation varies greatly across Europe, largely due to climatic factors and local husbandry practices (see above). To gain more insight into general factors explaining the variation in wolf-caused incidents, we thus focused on the combined incidents across all species.

Across the NUTS3 regions, only 2.1% of the total variation in the number of wolf-caused incidents was within regions across years, while 97.9% was across regions (*p* < 10^-15^). To explain that latter part, we conducted regression analyses on the average number of wolf-caused incidents across years within each NUTS3 region. As explanatory covariates we used 1) the total area occupied by wolves, either sporadically (*Spo-A*) or permanently (*Per-A*), 2) the area within regions occupied by wolves for the CORINE land cover classes Pastures (2.3.1, referred to as P-A), Broad-leaved forests (3.1.1, *BLF-A)*, Coniferous forest (3.1.2, CF-A), Mixed forests (3.1.3, *MF-A)* and Natural grasslands (3.2.1, NG-A), 3) whether or not wolves were present at their lowest extent during the 1950-1970s (*50ya*), 4) the degree of governmental support for prevention measures (Prev) and 5) the mean temperature of the coldest quarter (BIO11) as a predictor for grazing season length.

We first used a Poisson model with log-link function and log-transformed covariates to explain average annual incidents rounded to their nearest integer. Using all ten covariates, that model explained *R*^2^ = 63.2% of the total variation and was the best fitting model according to the Akaike Information Criterion (AIC) (Supplementary Table S.4) and significantly better than the next-best sub-model with fewer covariates (ΔAIC = 12.76, Burnham and Anderson, 2004). However, and while all covariates significantly correlated with average annual incidents counts (*p* < 10^-27^), the majority of the variance appeared to be explained by a rather small set of covariates (Figure 2). By itself, *NG-A*, for instance, explained *R*^2^ = 47.5%, which was more than 3/4 of the total variance explained by all covariates. Other covariates with high explanatory power included *Per-A* (*R*^2^ = 37.6%), *BLF-A* (*R*^2^ = 36.6%), *MF-A* (*R*^2^ = 22.9%), *CF-A* (*R*^2^ = 16.2%) and *50ya* (*R*^2^ = 10.1%), while the remaining had *R*^2^ < 10% and *BIO11* even only *R*^2^ = 0.4%. Allowing for two covariates, the best model included *NG-area* and *Per-A* and explained *R*^2^ = 57.1% of the total variance, or > 90% of the variance explained using all ten covariates. The best models using three covariates further included *BLF-A* and explained > 95% of the variance explained by all ten covariates, while five covariates were sufficient to explain > 99%. Notably, a model just including *Per-A* and *Spo-A* explained *R*^2^ = 37.7% and thus significantly less than *NG-A* alone (ΔAIC = 3243.18).

We obtained qualitatively similar results when using a Gaussian model on the raw covariates with power-transformed incident counts, for which we estimated the best transformation exponent to be *τ* = 0.54. In this case, the total variance explained by all covariates was *R*^2^ = 56.6%, while the best model according to AIC explained *R*^2^ = 56.0% using the seven covariates *Per-A, Spo-A, NG-A, CF-A, P-A* and *BIO11* (Supplementary Table S.4). There were, however, several models not significantly different from the best model (ΔAIC < 2.0), including a model using all covariates except *Prev*. Again, *NG-A* had the biggest individual explanatory power with *R*^2^ = 41.3%, matched 3/4 of the total variance explained by the best model and explained significantly more than the *R*^2^ = 16.1 of a model including *Per-A* and *Spo-A* (ΔAIC = 154.82). The covariates with the next highest explanatory power, however, differed in order from the Poisson model and included *BLF-A* (*R*^2^ = 21.4%), *Per-area* (*R*^2^ = 15.9%) and *50ya* (*R*^2^ = 13.4%). The remaining had *R*^2^ < 10%, again with *BIO11* explaining the least (*R*^2^ < 0.1%) and the only covariate not significantly correlated (*p* = 0.76).

To avoid spurious fitting, the results above were obtained after excluding the outlier region of Asturias, Spain, which had more than twice as many annual incidents than the next region and thus contributed disproportionate to the total variance. When including this region, however, results were qualitatively similar: *NG-A* explained the most variance under both a Poisson and Gaussian model, the best model with two covariates included additionally *BLF-A* in both cases, and the best model included all ten covariates in the Poisson case but fewer covariates in the Gaussian case (Supplementary Figure S.1, Supplementary Table S.5).

Residuals of the best models were significantly spatially autocorrelated (*I* = 13, 554.8 and *I* = 843.2 under the Poisson and Gaussian models without outlier, *p* < 10^-6^ in both cases), suggesting that some additional variance may be explained with landscape features or other spatial factors not included in our model. To test if our choice of covariates was lacking any additional major environmental effect readily available for all NUTS3 regions, we extended our models with altitude, latitude and all bioclimatic variables available from WorldClim. Under the Poisson model (without outlier), adding all these 20 additional covariates explained an additional 9.2% of the total variation (*R*^2^ = 72.4%). However, this is likely a result of over-fitting and difficult to interpret: when added to the base model of ten covariates, the most informative additional covariate (*BIO9*) explained a mere extra 1.3% of the total variation, and all others an extra 0.4% or less. Similar results were obtained under the Gaussian model, where the most informative covariate (also *BIO9*) explained an extra 0.3% only.

In contrast, several interaction terms among the ten chosen covariates appear meaningful. Under the Poisson model, 43 of the 45 possible interaction terms led to significantly better models (ΔAIC > 2.0) when added individually to the model containing all ten considered covariates. Of those, two explained more than an extra 4% of the total variation: *50ya* × *Spo-A* (*R*^2^ = 68.0%, *β* = 0.44) and *50ya* × *NG-A* (*R*^2^ = 67.5%, *β* = −0.43). Under the Gaussian model, 24 of the 45 possible interaction terms led to significantly better models, of which five explained an extra 4% or more of the total variation: *NG-A* × *P-A* (*R*^2^ = 66.0%, *β* = −0.00007), *NG-A* × *BIO11* (*R*^2^ = 62.2%, *β* = 0.0028), *NG-A* × *Prev* (*R*^2^ = 61.7%, *β* = 0.0016), *NG-A* × *50ya* (*R*^2^ = 61.0%, β = −0.0200) and *BLF-A* × *50ya* (*R*^2^ = 61.0%, *β* = −0.0099).

### 3.4. Number of wolf units correlated with incidents

For Germany, detailed information is available on the number of wolf units in each NUTS region for each of the three years studied here. Focusing on the NUTS3 regions for which livestock damage incidents, wolves or both were reported, the number of wolf units was significantly correlated with the number of reported incidents for each year (*p* < 0.001, *p* < 0.001 and p = 0.002, respectively) as well as for all years combined (*p* < 0.001). The magnitude of the correlation diminished over time, from ρ = 0.60 in 2018, to ρ = 0.49 and ρ = 0.39 in 2019 and 2020, respectively.

Interestingly, the average number of wolf units across the three years was a slightly worse predictor of the average number of reported incidents per region within Germany than the *Per-A* and *Spo-area* derived of a distribution map for 2012-2016 (*R*^2^ = 0.37 vs. *R*^2^ = 0.38 for the Poisson model and *R*^2^ = 0.37 vs. *R*^2^ = 0.44 for the Gaussian model).

### 3.5. Trend analysis

We estimated trends of wolf-caused incidents across the three years 2018, 2019 and 2020 for all geographic regions with at least one incident reported from at least two years, accounting for survey gaps and stochastic variation. At the continental scale, our analysis indicated with certainty that incidents were increasing 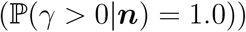 with an estimated rate of 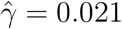 per year (posterior mode), translating into an 4.2% increase from 2018 to 2020. At smaller geographic scales (Figure 4), the pattern is rather heterogeneous: of the 320 NUTS3 regions with sufficient data (two years with damage incidents), we estimated a positive trend 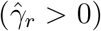 for 195 (61%) and a negative trend 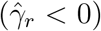 for 125 (39%), with posterior modes spanning from 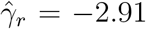 for NO074 (Troms og Finnmark, Norway) to 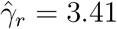 for FRC13 (Saône-et-Loire, France). Despite this heterogeneity, the 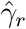 estimates, and thus the directionality of the trends, were spatially autocorrelated (*p* < 0.003), but not correlated with the recency of colonization, neither when using the historical wolf distribution *50ya* (*U* = 6250, *p* = 0.75) nor when comparing regions in the center of the wolf distribution (all neighboring regions had damages) to those at the frontier (bordering regions without damages, *U* = 1347, *p* = 0.24).

**Figure 4:**
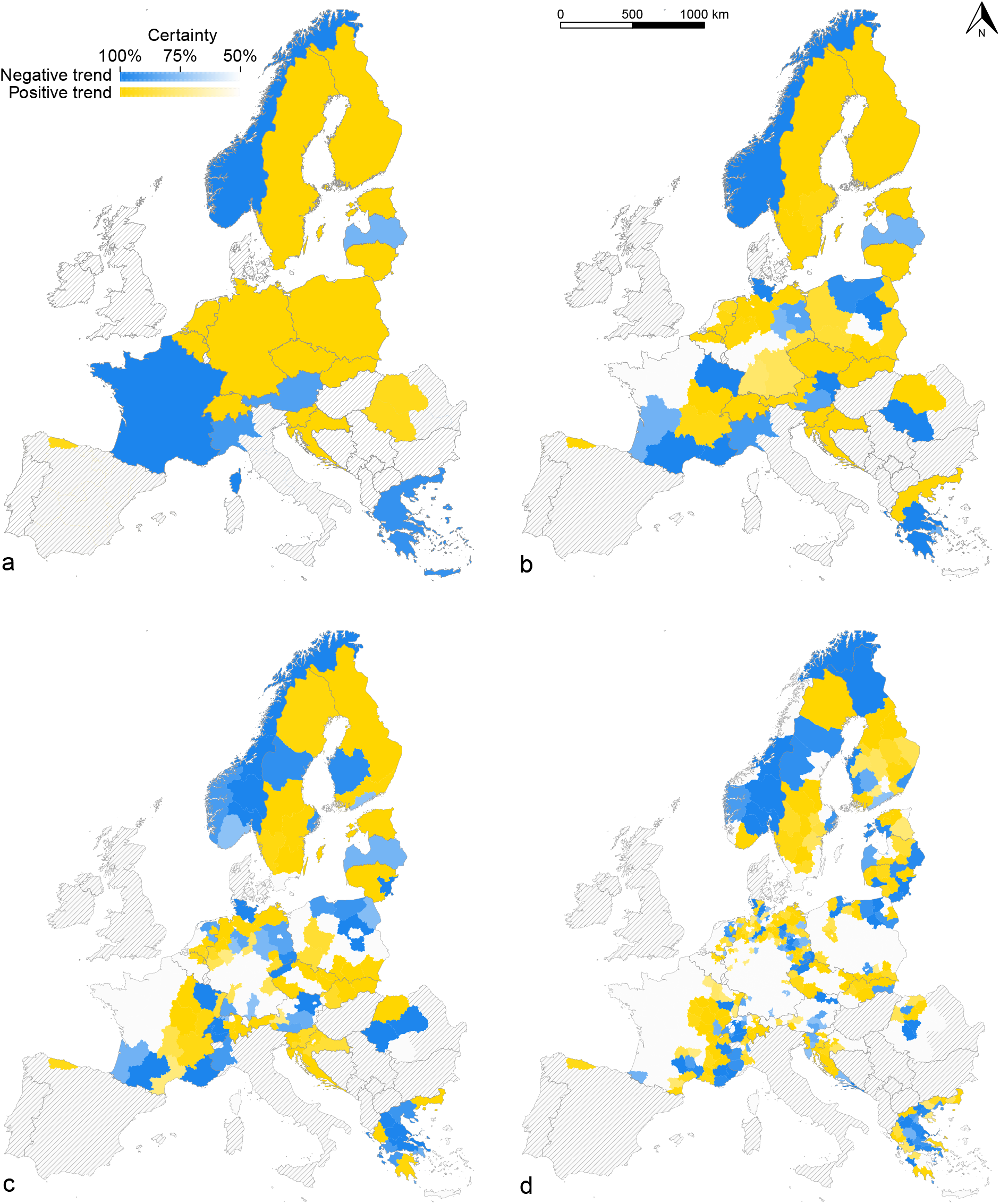
Wolf-caused livestock damage trends at the country (a), NUTS1 (b), NUTS2 (c) and NUTS3 (d) level. Each region was classified as having an increasing (yellow) or decreasing (blue) incident trend, depending on the posterior mode *γ_r_* < 0 for decreasing trends and *γ_r_* > 0 for increasing trends. Color saturation indicates uncertainty quantified as the posterior mass within the classified interval, ranging from solid (1.0) to white (≤ 0.5). Regions without reported incidents are shown as white, regions with no data reported are shaded in gray.

We also estimated trends individually for the most commonly affected species (sheep, goats, cattle and horses, Supplementary Figures S.2–S.5). Trends did not appear to be correlated between any pair of species at NUTS1, NUTS2 or NUTS3 level (p> 0.06 in all cases), likely because trends could be estimated only for a partially overlapping subset of regions for each species due to the lower number of incidents and a restricted geographic distribution of some species.

For Germany, we also estimated trends in the number of wolf units for each NUTS3 region with at least one wolf unit reported across the three years 2018, 2019 and 2020. The joint trend across all such regions revealed a rapidly growing population 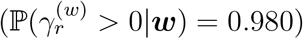 with 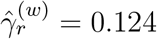 (posterior mode), corresponding to the growth rate of 13.2% per year. To confirm that growth rates decreased over time, we estimated them for all three-year intervals from 2006 to 2020 for which at least two NUTS3 regions had wolves in three years. Estimated annual growth rates (posterior modes) decreased from 65.8% for 2006-2008 to 33.6%, 32.2%, 30.9% and finally 13.2% for 2018-2020.

There was considerable regional variation, with 47 NUTS3 regions showing positive 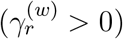 and 26 negative 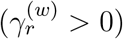 trends of wolf units for 2018-2020. We tested whether these trends predict trends in wolf-caused incidents, but did not find such a correlation at any NUTS level (*p* > 0.15 for Spearman correlations and *p* > 0.08 for Pearson correlations in all cases). The trends in wolf-caused incidents did not correlate with the time since wolves were first reported in a region (*p* > 0.16 at any NUTS level). Across Germany, however, wolf units 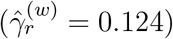 and damage incidents 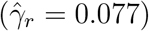 did not grow at significantly different rates 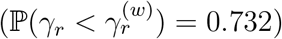.

## 4 Discussion

We here consolidated a large, incident-based data set on livestock damages caused by wolves across Europe in recent years. A total of 16 countries reported complete data for the years 2018, 2019 and 2020, and an additional five countries reported partial data. The majority of reported incidents involved a single livestock head and only very few involved more than ten individuals. In line with previous reports (Kaczensky, 1999; Bautista et al., 2019; Gervasi et al., 2021a), sheep were the most affected species, both in terms of incidents and affected individuals. There was, however, spatial variation reflecting the regional importance of different livestock species such as reindeer in Finland or horses in Spain. Interestingly, however, cattle suffered disproportionately many incidents in Greece, although Greece had the least amount of cattle per sheep in 2019 (according to EuroStat, Table apro-mt_ls) of all EU member states for which we received damage data.

We found considerable seasonal variation with incidents peaking in August and September. There was a clear north-south cline with a much smaller fraction of incidents reported during winter in northern compared to southern Europe, likely because in the north, livestock is kept indoors more often during these months. Finland showed a particular interesting seasonal pattern in the number of reported reindeer incidents, which are much higher in early than in late winter (Figure 3). This is largely due to the migration of wolves into Finnish reindeer husbandry areas in autumn, leading to many reported incidents, followed by their legal hunting later in winter, which reduces their numbers and consequently reported damage incidents.

### 4.1. Extensively used habitats favor wolf-caused damage incidents

The data also revealed substantial spatial variation in the number of incidents caused by wolves across Europe. A large part of that variation (>40%) is explained by a single environmental covariate: the area of natural grasslands of each region occupied by wolves. Up to about 60% of the total variation is explained by just a few additional covariates such as the area permanently occupied by wolves or the area of broad-leaved forest occupied by wolves, though their ranking varied with model choices. Regardless of these choices, however, a model only including the area permanently or sporadically occupied by wolves had a much poorer fit. Thus, areas with high numbers of incidents are not only qualified by the presence of wolves, but also through extensively cultivated habitats where wolves may be numerous and livestock more accessible or more difficult to protect (Rigg et al., 2011; Jedrzejewski et al., 2004; Treves et al., 2004).

Correlations for similar landscape features were recently reported from damages in Poland (Fedyń et al., 2022), but not from a recent, multi-national analysis on sheep incidents for which compensation was paid (Gervasi et al., 2021a). Based on that data, they reported an effect of historical continuity of wolf presence, but unlike our study found no correlation with any of the environmental variables tested. This apparent difference is likely due to three reasons: First, our data set encompasses many more NUTS3 regions (470 vs. 140) and hence has more statistical power to detect such correlations. Second, Gervasi et al. (2021a) modeled yearly damage counts for each region through year- and region-specific random effects. We decided against such a strategy and rather focused on average counts per region, because annual count data are not strictly independent due to, for instance, local husbandry practices and the same wolf individuals causing damage across multiple years. While region-specific random effects may capture that effect, they effectively deprive the model of identifying fixed effects. Third, we considered landscape features only within the areas predicted to be occupied by wolves, rather than on NUTS3 regions as a whole. Indeed, most NUTS3 regions are only partially occupied by wolves, potentially due to historical or political reasons not connected to the landscape features of the entire region. Consequently, and for the Poisson and Gaussian cases respectively, the best models using the restricted areas explained 2.6% and 10.0% more of the total variance than the best models using the landscape features of the entire NUTS3 regions. On top of such land cover metrics, the historical continuity of wolf presence then appeared to have significant but limited explanatory power.

While our models captured a large part of the between-region variation on wolf-causes damage incidents, about 40% of it remained unexplained even with our best fitting model. Part of that variance may be due to variation in reporting rates between regions, rather than variation in damage incidents themselves (Gervasi et al., 2021a). Also, incident numbers may be inherently stochastic, especially if wolf numbers are low such that a single problem individual can cause a spike in incidents for a short time. In the NUTS2 region AT12 (Niederösterreich, Austria), for instance, most of the 19 incidents in 2018 could be attributed to a single female wolf. This individual was no longer active in 2019 and 2020, when incidents dropped to two and only one, respectively. In our data set, however, only 2.1% of the total variation was across years.

Since residuals appeared to be geographically clustered such that too many (or too few) incidents were often predicted for multiple neighboring regions, there likely exist additional covariates predictive of incident numbers. Promising such covariates include the densities of wild prey species (Meriggi and Lovari, 1996), with wolves both more likely to switch to livestock in areas of low densities (Kaartinen et al., 2009a), yet particularly attracted to areas with hight densities (Treves et al., 2004), increasing damage incidents in both. While estimates of wild prey densities are currently not available at the continental scale, they may themselves be explained through bioclimatic and land-use variables. When adding all bioclimatic variables of WorldClim as well as altitude and latitude, an extra almost 10% of the variation in average annual incidents counts could indeed be explained, but no single of those additional covariates appeared to explain sufficient extra variation to warrant their discussion.

Interactions among covariates had larger individual contributions, but given their large number, significant testing becomes difficult. It is nonetheless interesting to note that many of the interactions that explained several extra percent of the total variation involved the historical continuity of wolves, always with negative coefficients, suggesting that the main environmental covariates appear to underestimate wolf-caused incidents for regions only recently occupied by wolves. This is in good agreement with the thought that incidents often spike in areas recently colonized by wolves (Trouwborst, 2018; Marucco and Boitani, 2012; Dalmasso et al., 2011; van Eeden et al., 2018; Gervasi et al., 2021a), as measures preventing livestock damages may have been abandoned in the absence of large carnivores (Kaczensky et al., 2021; Linnell et al., 1996). While the historical continuity of wolf presence since the 1950-1970s is unlikely to capture this effect in full, only limited data is available to test for more recent effects. At the continental scale, for instance, the most recent estimate of wolf presence dates to the period of 2012-2016 Kaczensky et al. (2021), potentially already outdated at the frontier of the ongoing recolonization of Europe by wolves as we received reports of wolf-caused incidents for 78 NUTS3 regions presumably without any wolves present. The very low number of average annual incidents reported from these regions (median of 1.0), however, remained well explained by zero-value area covariates. And for Germany, for which more detailed and up-to-date data on the presence of wolves is available and for which 23 NUTS3 presumed wold-free regions reported incidents, the wolf area covariates derived from the outdated distribution maps turned out to be even better predictors of the number of average annual incidents than the actual number of wolf units present.

While wolves were not present in most regions within Germany about ten years ago, they have since made a successful comeback (BIJ12 et al., 2022): While Reinhardt et al. (2019) reported an annual growth rate of 36% for the period 2000-2015, it appears that the growth has been slowing down steadily to 14% for the three years considered here. Along with wolves, livestock damage incidents have increased at comparable rates at the larger scale, but the connection between the number of wolves and the number of reported incidents appears complex at local scales: first, we did not find any correlation between trends in the number of wolf units and trends in the number of damage incidents on the NUTS3 level. Thus, a growing wolf population does not seem to imply a growing number of incidents, a finding previously reported for sheep lost to wolves in Germany (Khorozyan and Heurich, 2022). Second, and while the number of wolves in a region seems to correlate well with the number of damage incidents, this correlation appears to diminish over time, in line with a lack of correlation between these trends. Thus, with wolves establishing themselves in more regions at higher numbers, the relationship between wolves and damage incidents becomes more complex.

### 4.2. Lack of data on the effectiveness of prevention measures

It may seem tempting to interpret these findings as evidence that a renewed presence of wolves leads to a more wide-spread adoption of protective measures, or at least that the relationship between damage incidents and wolf presence is modulated by variation in the adoption of such measures. However, the dataset gathered here is not ideal to directly test such hypotheses, nor hypotheses about the effectiveness of different prevention measures, as we lack information on their use. To assess damage prevention effectiveness, the frequency of damage incidents should be contrasted between sites differently protected. Authorities, however, tend to only record the prevention measure in place at the time of an incident, if at all, which does not allow conclusions on their effectiveness. In our dataset, for instance, the most frequently reported prevention measure was *electric fence*, most likely not because it was ineffective, but because it was very commonly used.

Despite the large amount of public money spent on supporting prevention measures, a large-scale randomized control trial on the effectiveness of different prevention measures is still crucially missing. As an alternative, the data reported here could be intersected with geo-referenced data on the use of prevention measures and accurate data on the presence of wolves, provided that the spatial auto-correlation in the use of prevention measures can be accounted for. To the best of our knowledge, such data is currently not available at a larger geographic scale. We thus strongly encourage authorities to collect and share such information in the future, ideally using specific categories that distinguish among the diverse set of measures treated under the same broad categories here. We note, however, that unbiased collection of such information may prove difficult if livestock owners fear consequences for improperly implemented measures.

### 4.3. Trends in damage incidents as indicators for human-wolf conflicts

By estimating trends in damage incidents, however, the data collected here provides useful indicators on the intensity of the conflict between wolves and livestock owners. These trends are comparable at any geographic level and robust to variation in sampling effort and data collection across regions, in contrast to analyses on raw incidents counts (Gervasi et al., 2021a). With wolf populations in Europe generally growing and expanding their range, decreasing trends indicate a reduction in conflicts, for instance through the adoption of additional prevention measures, other changes in husbandry practices, or the successful management of problematic individual wolves. Increasing damage incident trends, in contrast, identify regions where current damage prevention and mitigation seem insufficient and thus require additional attention. Nearly stable numbers of incidents, finally, indicate conflicts at equilibrium, either because damage prevention practices have been put in place successfully, because wolf numbers plateaued, or both.

Across the years 2018-2020 studied here, the majority of regions showed an increasing trend, reflecting the growth of the European wolf population and the need for conflict mitigation. In Germany, for which we have detailed information on the distribution of wolves, for instance, the number of wolf units and damage incidents grew at comparable rates. While we lack information on the growth of wolf populations at the continental scale, regional trends in damage incidents perfectly illustrate the small-scale nature of livestock damage incidents and their mitigation: first, we estimated negative trends for 39% of all NUTS3 regions, although wolf populations unlikely shrank in any of these.

Second, and while the inferred trends were spatially auto-correlated with incidents either increasing, decreasing or showing no trend in multiple neighboring regions, trends did not point uniformly in one direction for any country, neither at NUTS3 or NUTS2 level. Rather, there are regions in each country that are currently affected by an increasing number of damage incidents. In contrast to a recent study on Italy (Gervasi et al., 2021b), we did not find trends to systematically differ between regions with established wolf populations compared to those more recently colonized. Third, trends inferred individually for each of the most commonly affected species were not correlated. While species-specific trends may serve as useful indicators for the conflict between wolves and owners of a particular livestock species, the observed variation in trends may also reflect the limited number of incidents per species and limited geographic overlap in their use. Of the 272 NUTS3 regions for which we could infer trends for the most commonly affected species (sheep), only for 119 NUTS3 regions (43.8%) trends could also be inferred for the second most commonly affected species (cattle). This overlap is further reduced to 5 NUTS3 regions (1.8%) when comparing sheep and reindeer. More importantly, however, trends were inferred here over just three years, and may hence be subjected to random fluctuations not entirely captured by the Poisson assumption, particularly if wolf numbers are low.

### 4.4. Conclusions

We here report the largest data set on wolf-caused damage incidents across Europe to date. This data set revealed substantial spatial variation in damage densities across Europe, of which more than half could be explained by environmental covariates and in particular by the area of natural grasslands and broad-leaved forests occupied by wolves. The data set further supported the notion that regions recently colonized by wolves experience particularly high numbers of incidents, although the relationship between a growing wolf population and incidents appeared rather complex and modulated by regional or even local factors. We argue that trends in damage incidents provide promising insights into the intensity of the localized state of the conflict between wolves and livestock owners, with decreasing damage incidents despite growing wolf populations identifying regions particularly successful at damage prevention, while rapidly increasing damage incidents identifying regions where current damage prevention is largely insufficient. We estimated that damage incidents were currently growing in about 60% of all NUTS3 regions studied and therefore invite regional and national authorities to continue integrating their damage data into accessible data bases, and to use trends over longer time periods, at larger geographic scales, or both, to effectively monitor and help mitigate human-carnivore conflicts across Europe.

## Supporting information

All Supplementary material

## Data availability statement

The data that support the findings of this study are openly available from Zenodo at https://doi.org/10.5281/zenodo.6821814.

## Acknowledgments

We would like to thank the following people for their assistance with the collection of our data: Anna Maria Rodekirchen, Barbara Burčul, Benedikt Gehr, Cedric Claude, Daniela Hilfiker, Magdalena Rusiłowicz, Peter Jaxgard, Piotr Chmielewski, Romana Uhrinová, Tõnu Talvi.

## Supporting Information

### S1. Supplementary figures

**Figure S.1:**
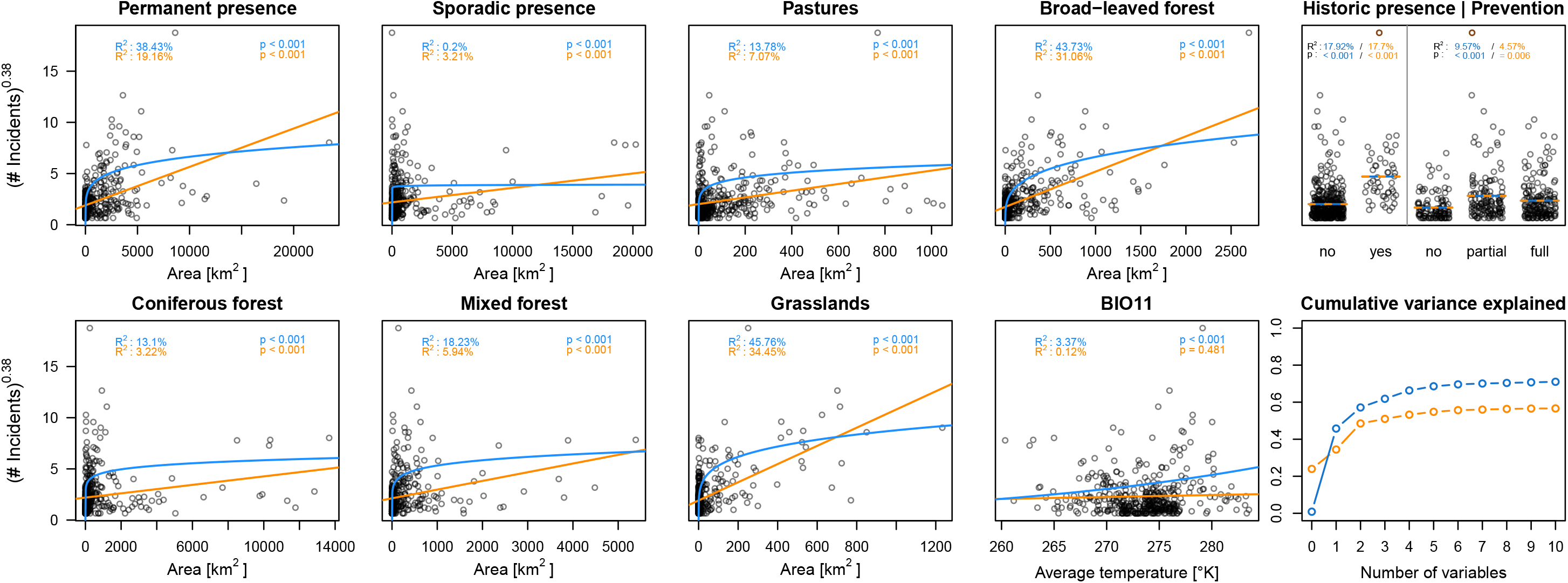
Correlation of ten environmental covariates and the number of reported wolf-caused livestock damage incidents in our data set, including the outlier region of Asturias (ES120). The Poisson model is shown in blue, the Gaussian model is shown in orange. For visualization purposes, both models are plotted using the same power-transformation on the incidents.

**Figure S.2:**
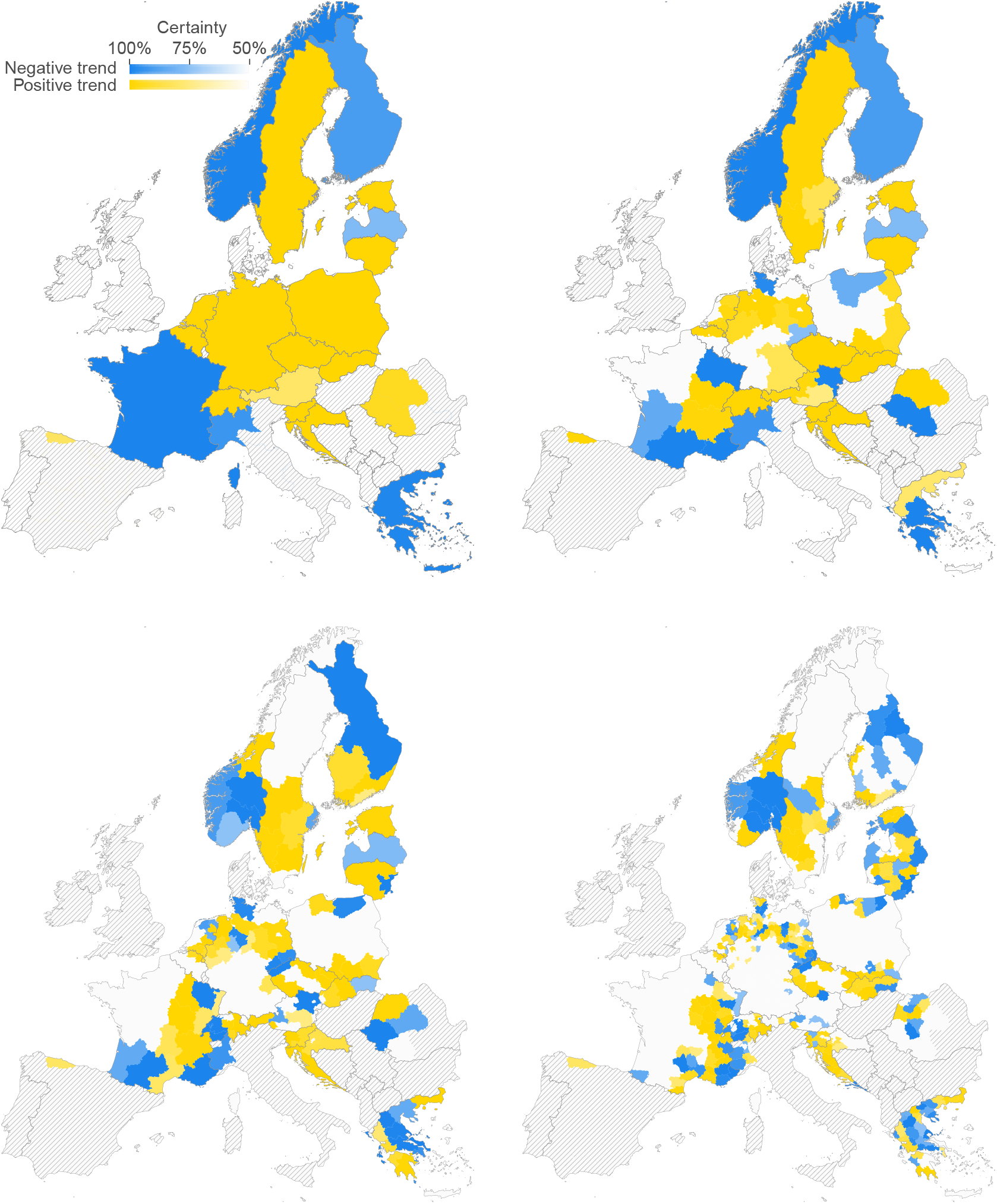
Wolf-caused livestock damage trends for sheep at the country, NUTS1, NUTS2 and NUTS3 level (top left to bottom right). Each region was classified as having an increasing (yellow) or decreasing (blue) incident trend, depending on the posterior mode *γ_r_* < 0 for decreasing trends and *γ_r_* > 0 for increasing trends. Color saturation indicates uncertainty quantified as the posterior mass within the classified interval, ranging from solid (1.0) to white (≤ 0.5). Regions without reported incidents are shown as white, regions with no data reported are shaded in gray.

**Figure S.3:**
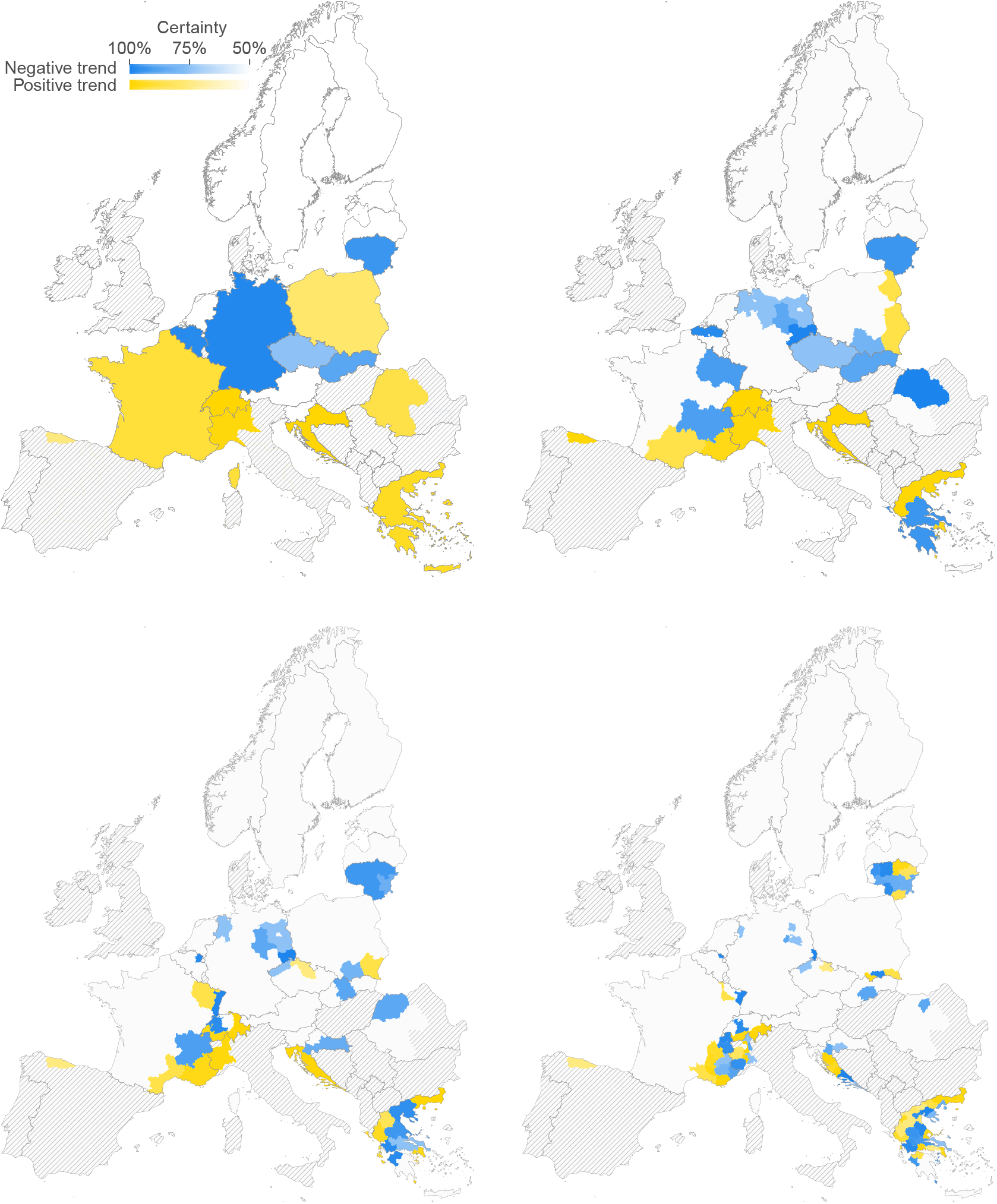
Wolf-caused livestock damage trends for goats at the country, NUTS1, NUTS2 and NUTS3 level (top left to bottom right). Each region was classified as having an increasing (yellow) or decreasing (blue) incident trend, depending on the posterior mode *γ_r_* < 0 for decreasing trends and *γ_r_* > 0 for increasing trends. Color saturation indicates uncertainty quantified as the posterior mass within the classified interval, ranging from solid (1.0) to white (≤ 0.5). Regions without reported incidents are shown as white, regions with no data reported are shaded in gray.

**Figure S.4:**
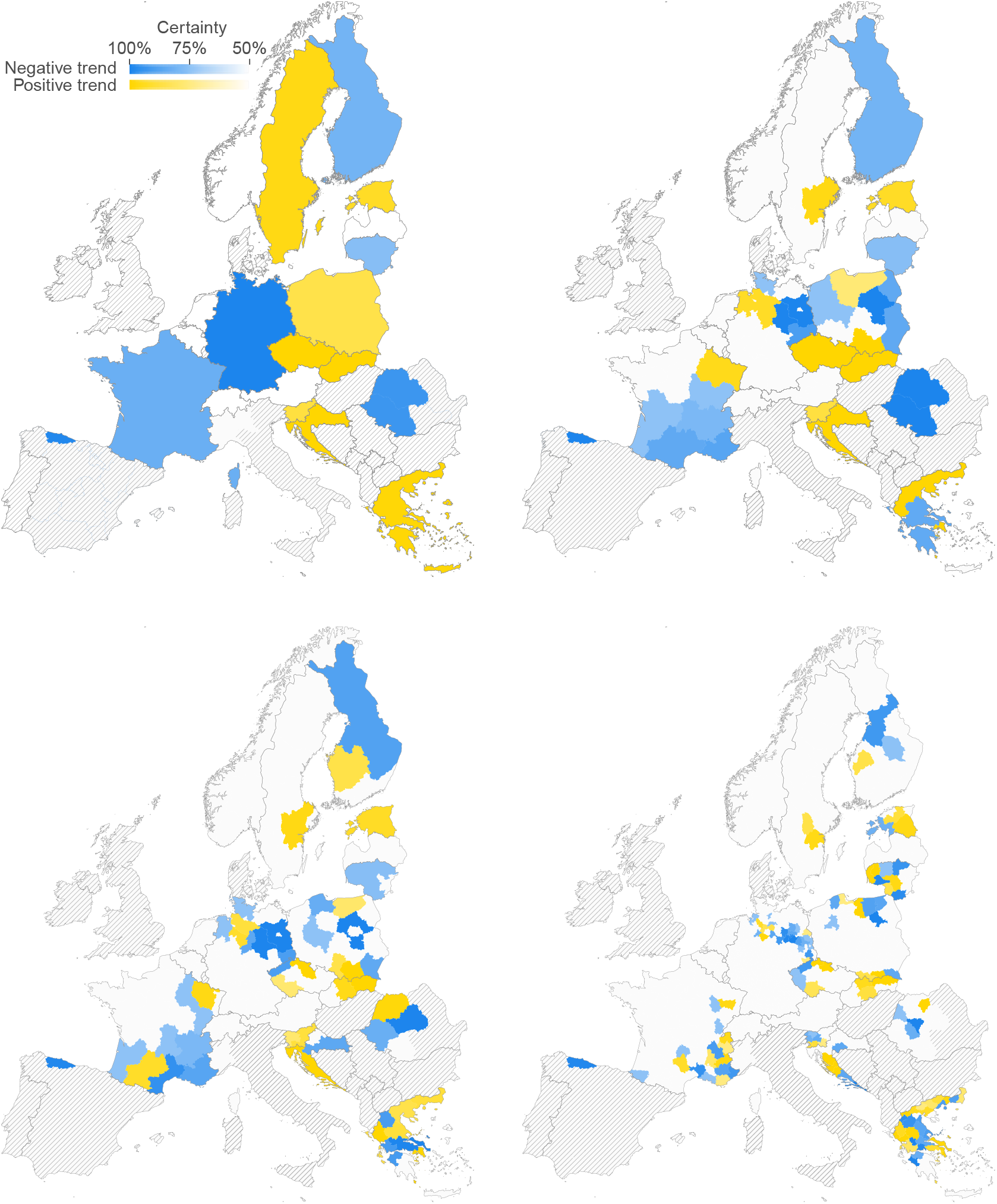
Wolf-caused livestock damage trends for cattle at the country, NUTS1, NUTS2 and NUTS3 level (top left to bottom right). Each region was classified as having an increasing (yellow) or decreasing (blue) incident trend, depending on the posterior mode *γ_r_* < 0 for decreasing trends and *γ_r_* > 0 for increasing trends. Color saturation indicates uncertainty quantified as the posterior mass within the classified interval, ranging from solid (1.0) to white (≤ 0.5). Regions without reported incidents are shown as white, regions with no data reported are shaded in gray.

**Figure S.5:**
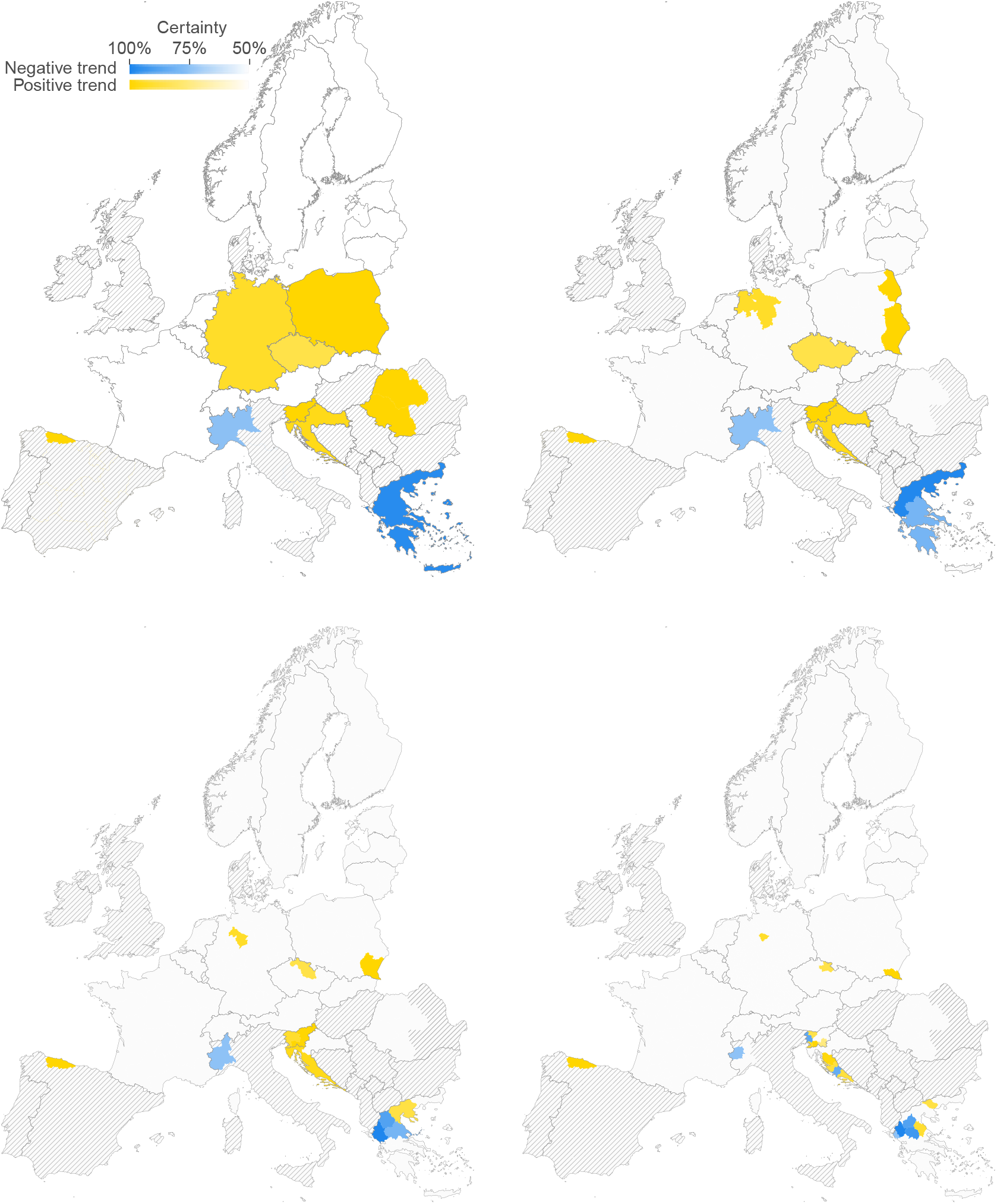
Wolf-caused livestock damage trends for horses at the country, NUTS1, NUTS2 and NUTS3 level (top left to bottom right). Each region was classified as having an increasing (yellow) or decreasing (blue) incident trend, depending on the posterior mode *γ_r_* < 0 for decreasing trends and *γ_r_* > 0 for increasing trends. Color saturation indicates uncertainty quantified as the posterior mass within the classified interval, ranging from solid (1.0) to white (≤ 0.5). Regions without reported incidents are shown as white, regions with no data reported are shaded in gray.

### S2. Supplementary tables

**Table S.1:**
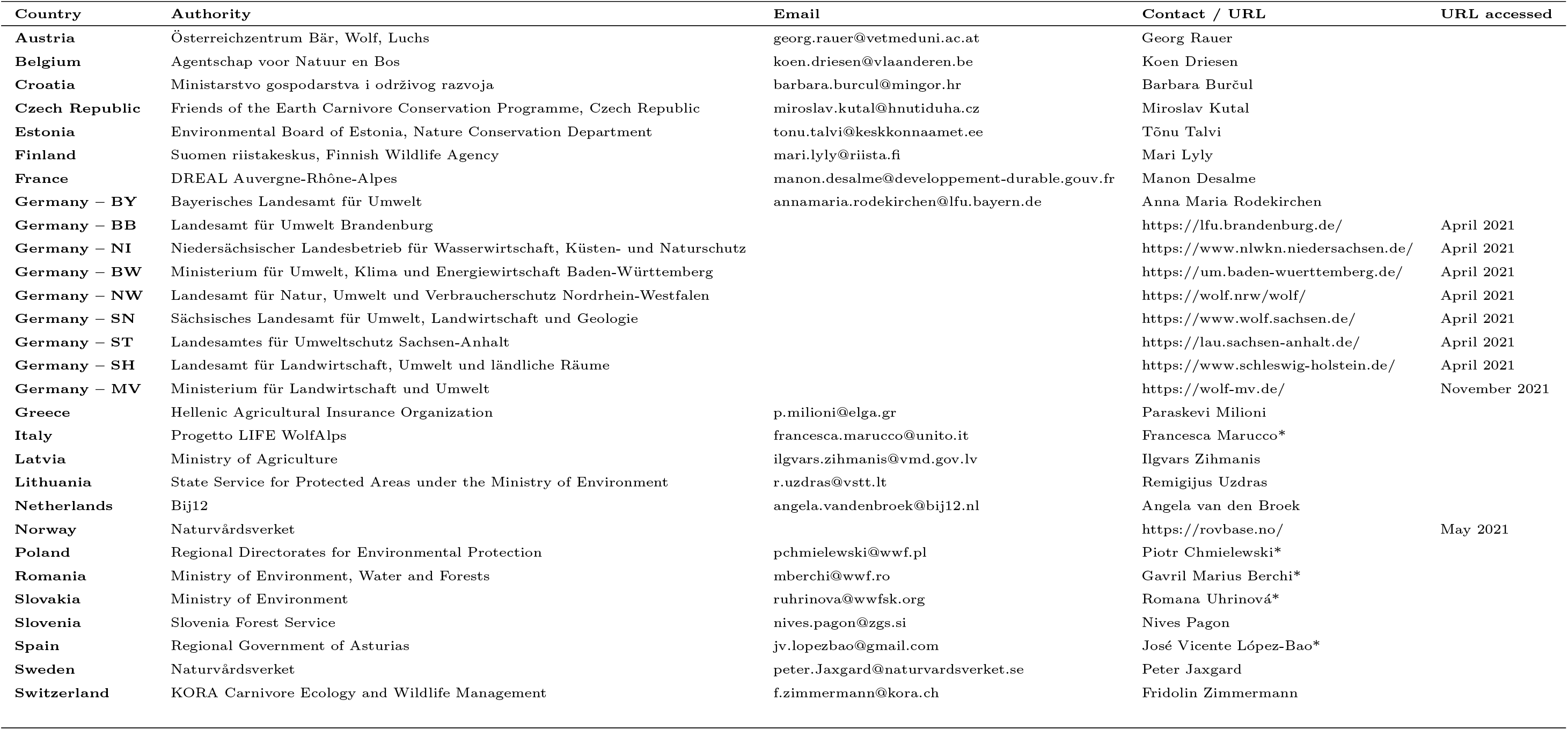
Sources of our livestock damage data. A * next to the contact name indicates that this person mediated the contact between us and the local authorities or this person was already collaborating with authorities and had permit to work with livestock depredation data.

**Table S.2:**
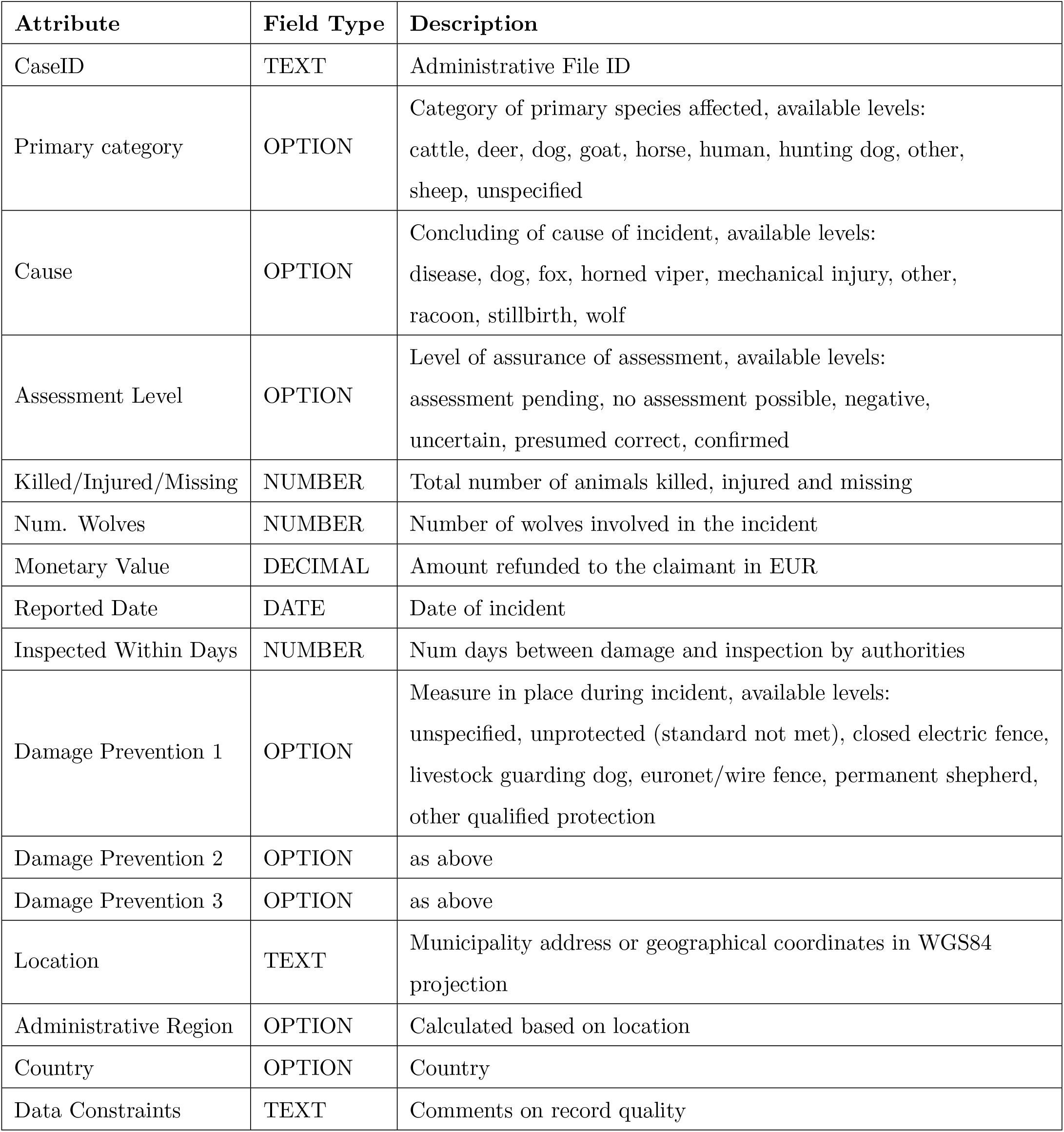
Livestock damage survey template.

**Table S.3:**
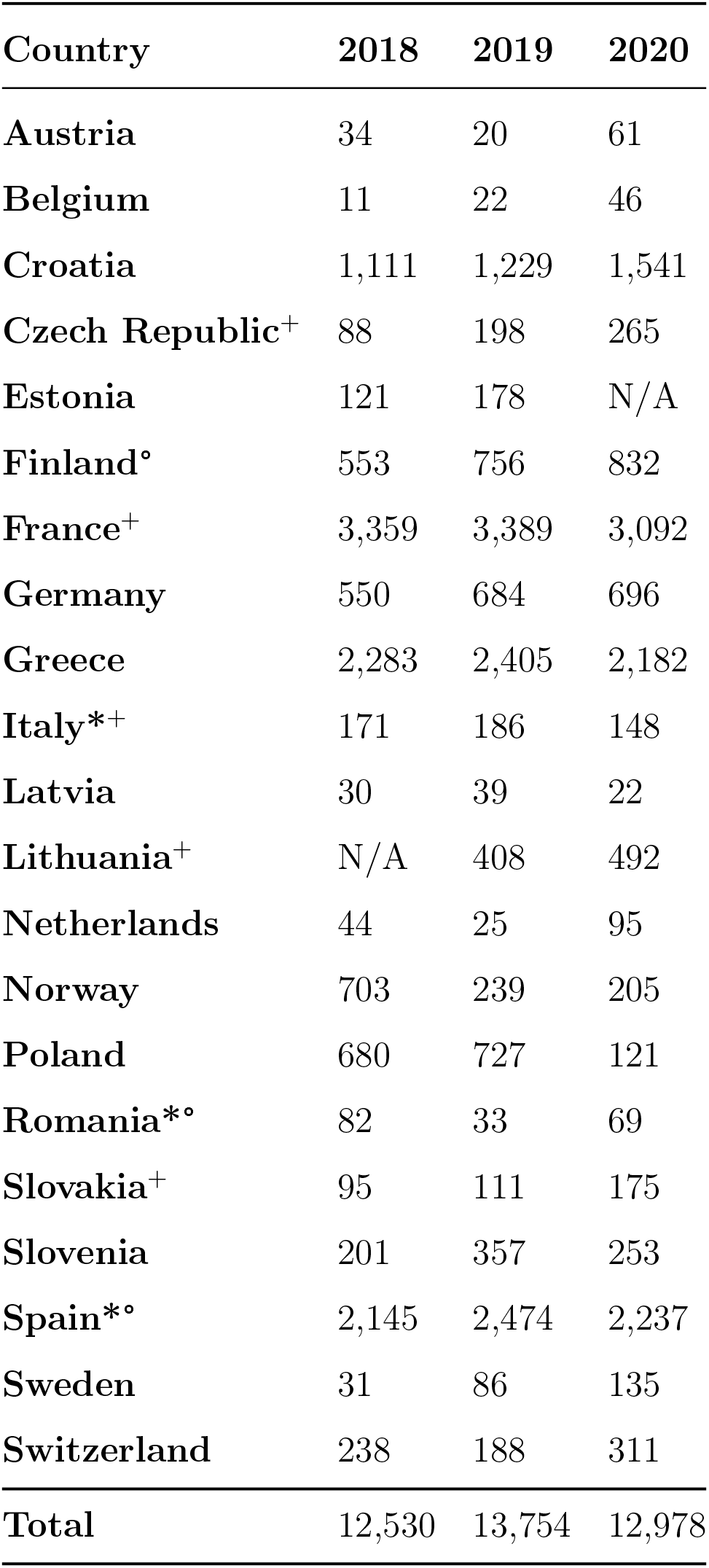
Number of wolf-caused livestock damage incidents per year and per country (see main text). Countries marked with a * reported data only for a subset of the provinces with documented wolf presence. Countries marked with a ^+^ did not report an assessment level and we therefore analyzed all submitted records. For countries marked with a *°*, we analyzed compensated incidents.

**Table S.4:**
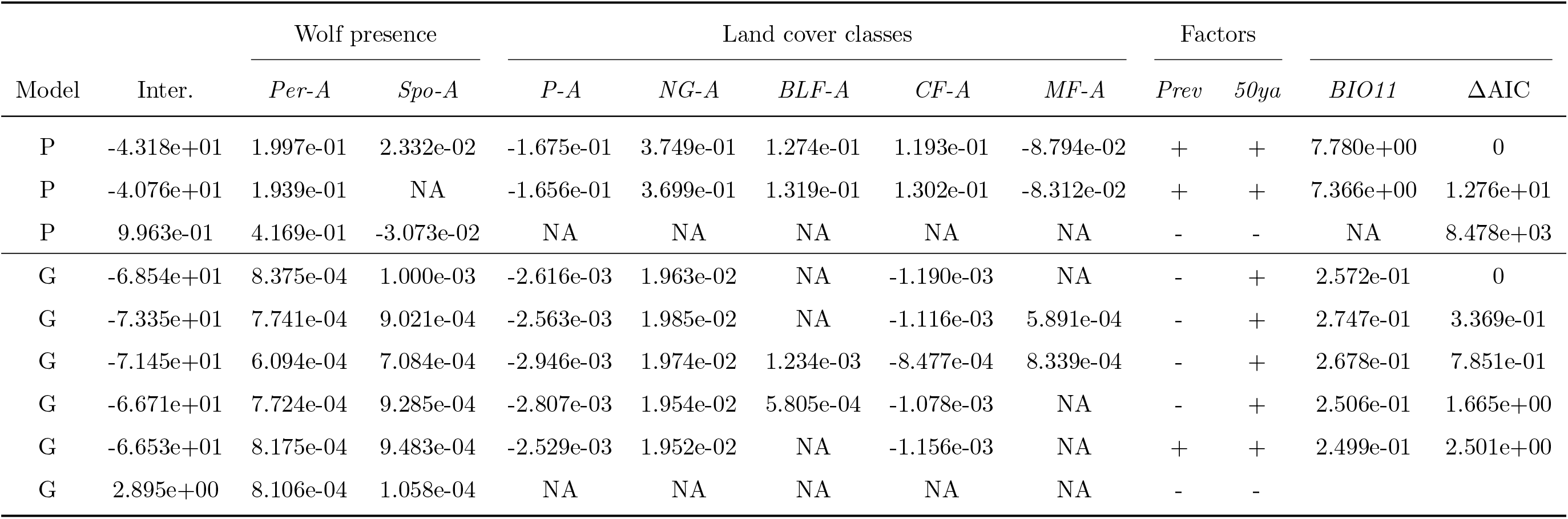
Generalized linear models fitted to the average number of wolf-caused incidents excluding the outlier region of Asturias (ES120). The table shows the models with the best fit (delta AIC < 2.0) as well as the model including wolf presence only. The first three rows correspond to the Poisson model (P), for which all covariates except the factors were log-transformed. The lower six rows correspond to the Gaussian model (G), for which the average number of incident counts were power-transformed (see main text).

**Table S.5:**
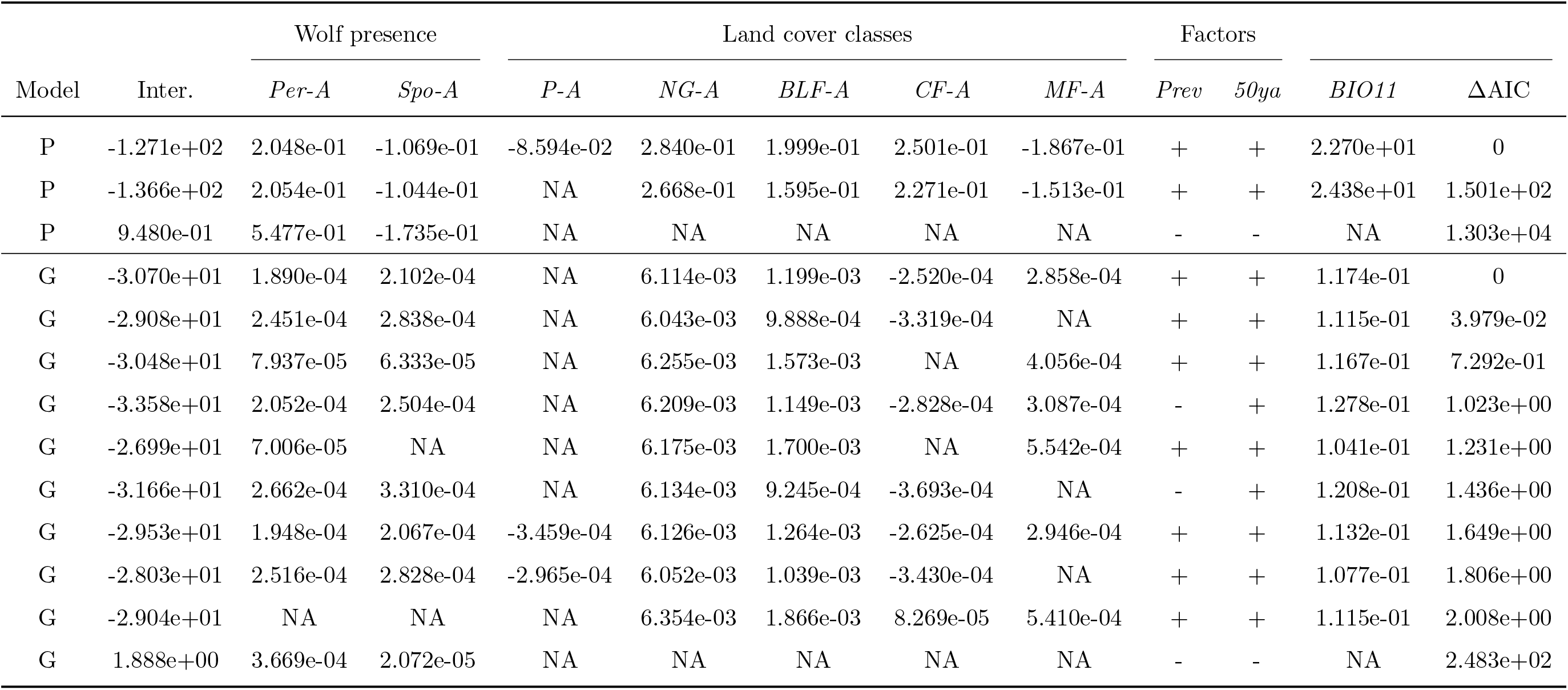
Generalized linear models fitted to the average number of wolf-caused incidents inculding the outlier region of Asturias (ES120). The table shows the models with the best fit (delta AIC < 2.0) as well as the model including wolf presence only. The first three rows correspond to the Poisson model (P), for which all covariates except the factors were log-transformed. The lower ten rows correspond to the Gaussian model (G), for which the average number of incident counts were power-transformed (see main text).

