## Supplementary material for "The spatial distribution and temporal trends of livestock damages caused by wolves in Europe": All Supplementary material

—

### Supporting Information

#### S1. Supplementary figures

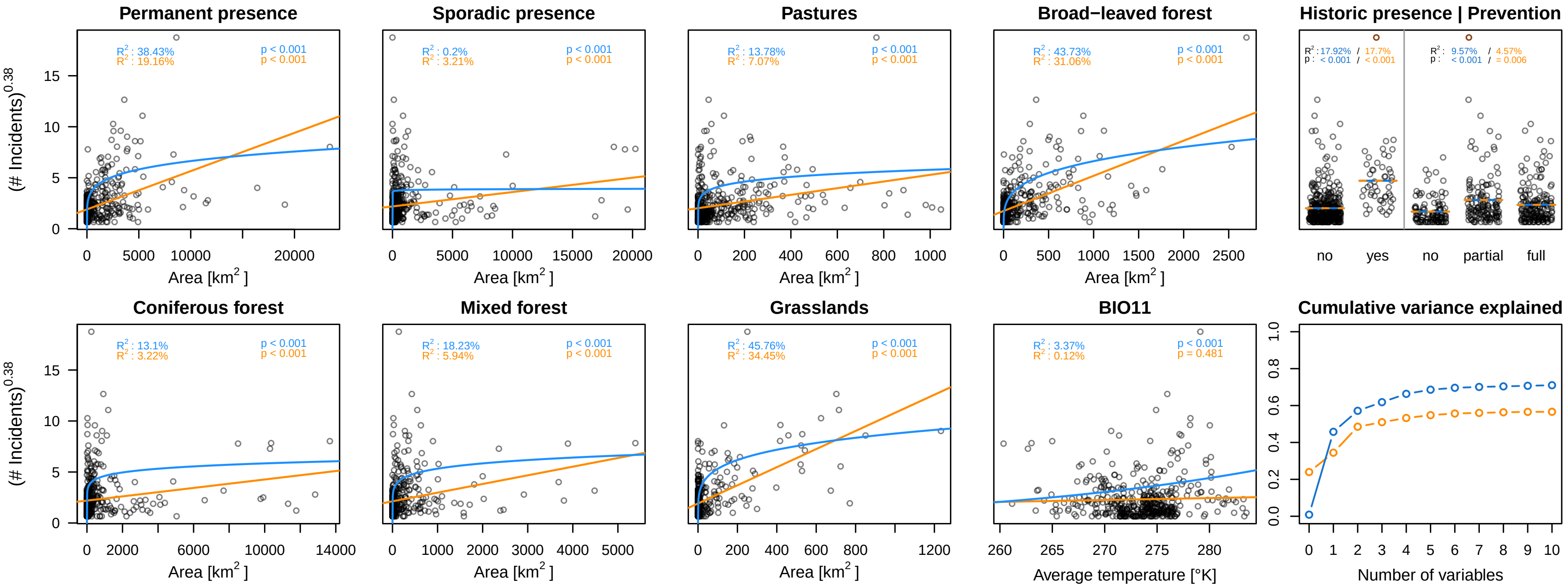


Figure S.1: Correlation of ten environmental covariates and the number of reported wolf-caused livestock damage incidents in our data set, including the outlier region of Asturias (ES120). The Poisson model is shown in blue, the Gaussian model is shown in orange. For visualization purposes, both models are plotted using the same power-transformation on the incidents.

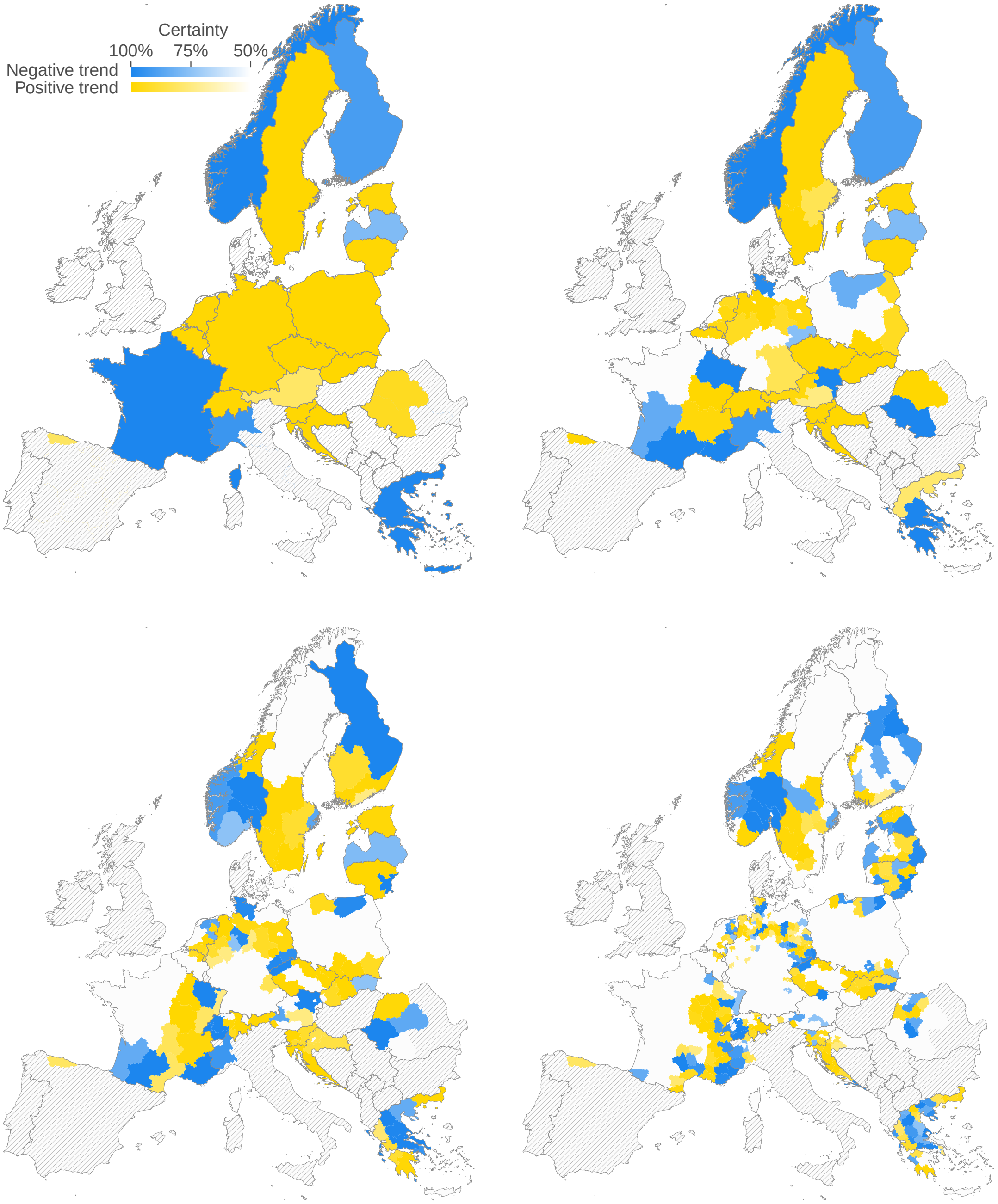


Figure S.2: Wolf-caused livestock damage trends for sheep at the country, NUTS1, NUTS2 and NUTS3 level (top left to bottom right). Each region was classified as having an increasing (yellow) or decreasing (blue) incident trend, depending on the posterior mode $\gamma_r < 0$ for decreasing trends and $\gamma_r > 0$ for increasing trends. Color saturation indicates uncertainty quantified as the posterior mass within the classified interval, ranging from solid (1.0) to white ($\leq 0.5$). Regions without reported incidents are shown as white, regions with no data reported are shaded in gray.

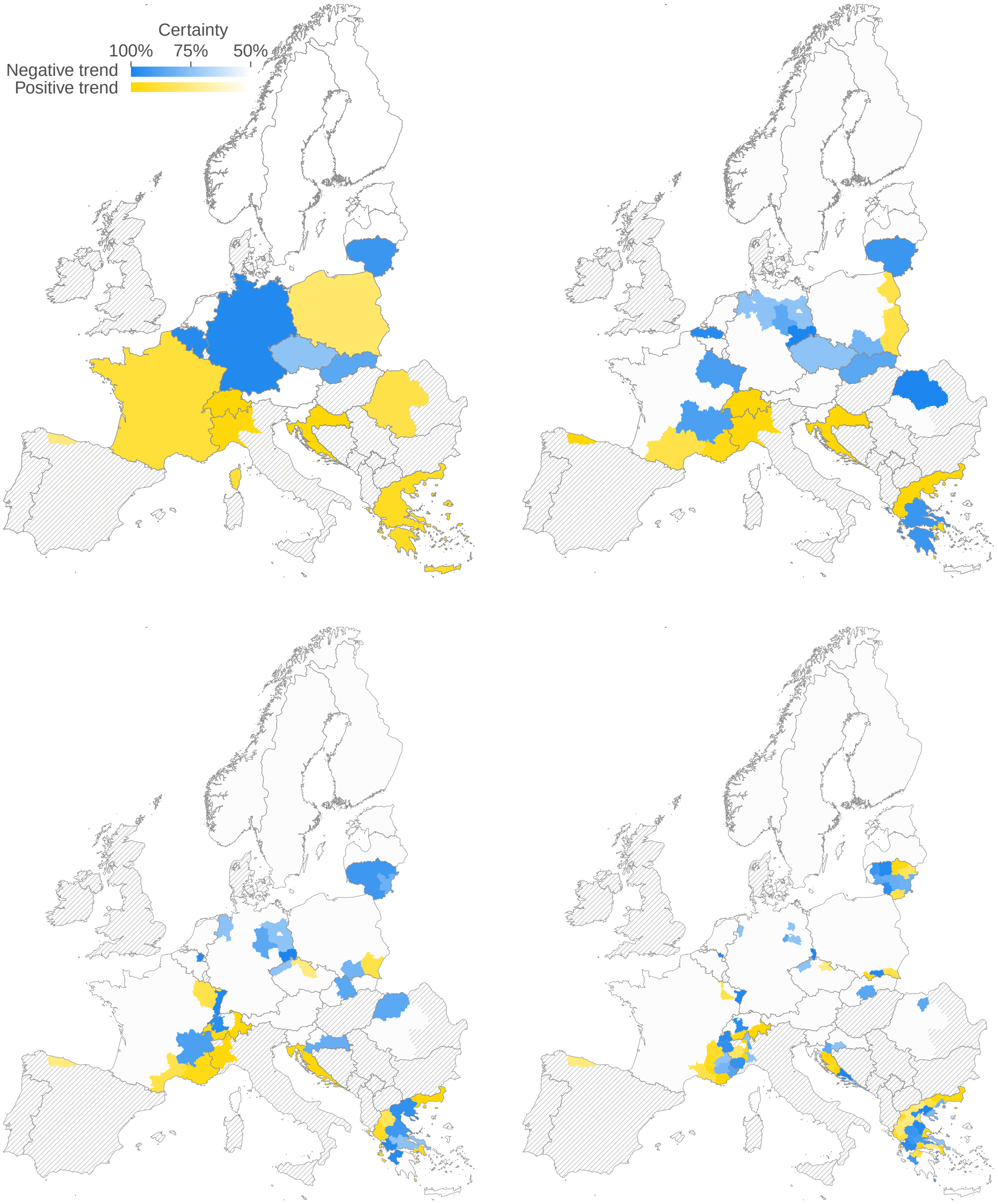


Figure S.3: Wolf-caused livestock damage trends for goats at the country, NUTS1, NUTS2 and NUTS3 level (top left to bottom right). Each region was classified as having an increasing (yellow) or decreasing (blue) incident trend, depending on the posterior mode $\gamma_r < 0$ for decreasing trends and $\gamma_r > 0$ for increasing trends. Color saturation indicates uncertainty quantified as the posterior mass within the classified interval, ranging from solid (1.0) to white ($\leq 0.5$). Regions without reported incidents are shown as white, regions with no data reported are shaded in gray.

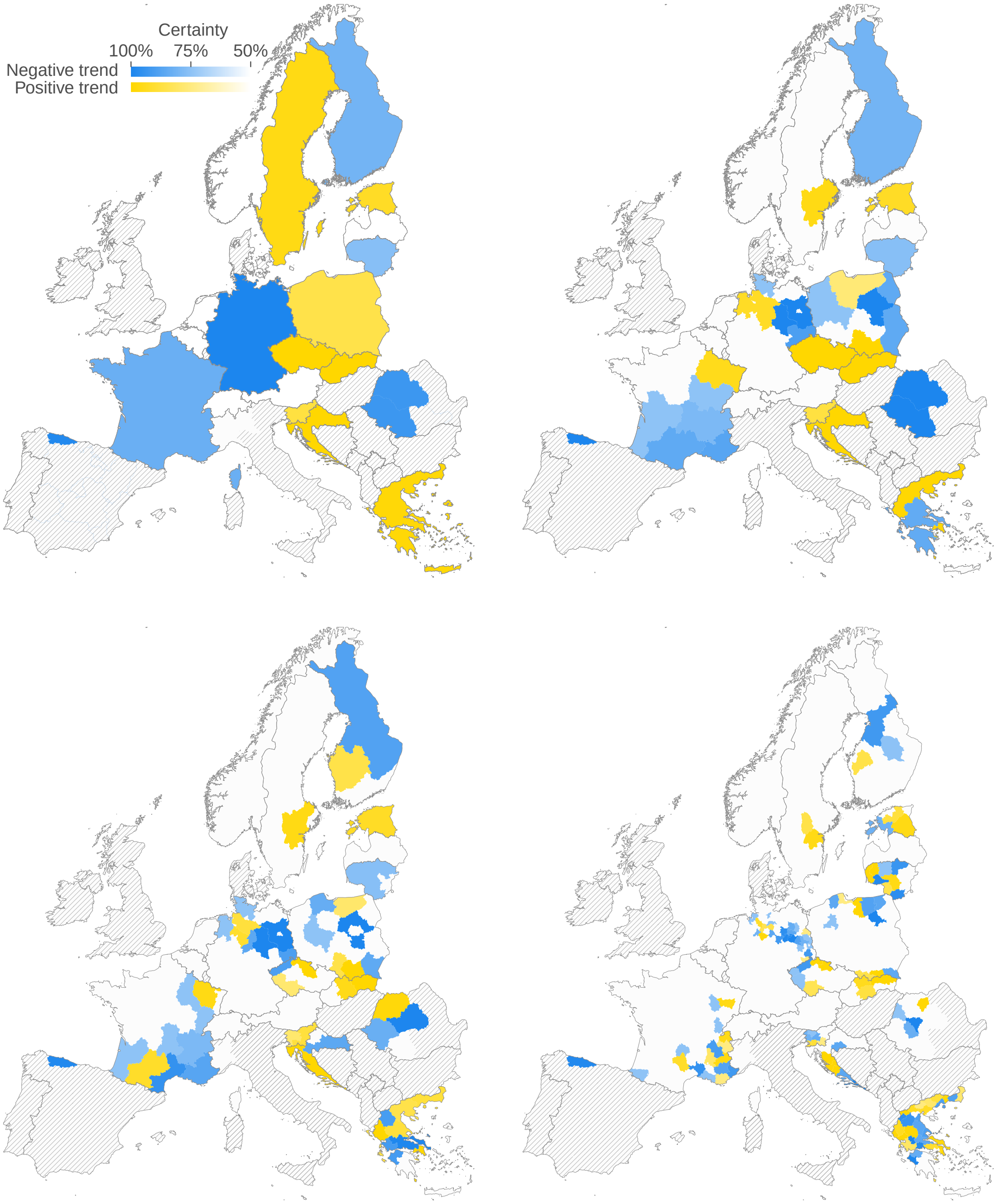


Figure S.4: Wolf-caused livestock damage trends for cattle at the country, NUTS1, NUTS2 and NUTS3 level (top left to bottom right). Each region was classified as having an increasing (yellow) or decreasing (blue) incident trend, depending on the posterior mode $\gamma_r < 0$ for decreasing trends and $\gamma_r > 0$ for increasing trends. Color saturation indicates uncertainty quantified as the posterior mass within the classified interval, ranging from solid (1.0) to white ($\leq 0.5$). Regions without reported incidents are shown as white, regions with no data reported are shaded in gray.

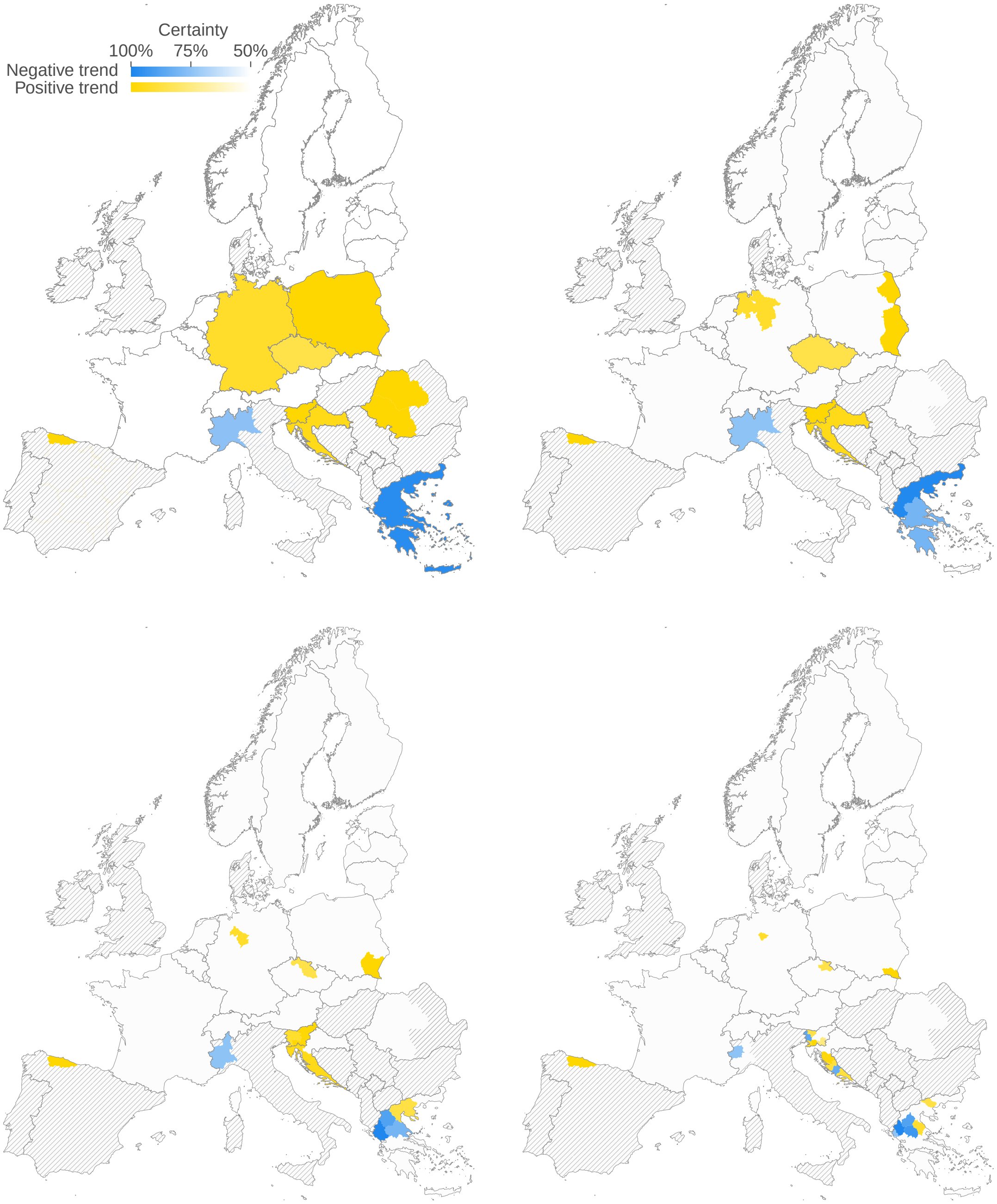


Figure S.5: Wolf-caused livestock damage trends for horses at the country, NUTS1, NUTS2 and NUTS3 level (top left to bottom right). Each region was classified as having an increasing (yellow) or decreasing (blue) incident trend, depending on the posterior mode $\gamma_r < 0$ for decreasing trends and $\gamma_r > 0$ for increasing trends. Color saturation indicates uncertainty quantified as the posterior mass within the classified interval, ranging from solid (1.0) to white ($\leq 0.5$). Regions without reported incidents are shown as white, regions with no data reported are shaded in gray.

#### S2. Supplementary tables

Table S.1: Sources of our livestock damage data. A * next to the contact name indicates that this person mediated the contact between us and the local authorities or this person was already collaborating with authorities and had permit to work with livestock depredation data.

| Country | Authority | Email | Contact / URL | URL accessed |
|---|---|---|---|---|
| **Austria** | Österreichzentrum Bär, Wolf, Luchs | | Georg Rauer | |
| **Belgium** | Agentschap voor Natuur en Bos | | Koen Driesen | |
| **Croatia** | Ministarstvo gospodarstva i održivog razvoja | | Barbara Burčul | |
| **Czech Republic** | Friends of the Earth Carnivore Conservation Programme, Czech Republic | | Miroslav Kutal | |
| **Estonia** | Environmental Board of Estonia, Nature Conservation Department | | Tõnu Talvi | |
| **Finland** | Suomen riistakeskus, Finnish Wildlife Agency | | Mari Lyly | |
| **France** | DREAL Auvergne-Rhône-Alpes | | Manon Desalme | |
| **Germany – BY** | Bayerisches Landesamt für Umwelt | | Anna Maria Rodekirchen | |
| **Germany – BB** | Landesamt für Umwelt Brandenburg | | https://lfu.brandenburg.de/ | April 2021 |
| **Germany – NI** | Niedersächsischer Landesbetrieb für Wasserwirtschaft, Küsten- und Naturschutz | | https://www.nlwkn.niedersachsen.de/ | April 2021 |
| **Germany – BW** | Ministerium für Umwelt, Klima und Energiewirtschaft Baden-Württemberg | | https://um.baden-wuerttemberg.de/ | April 2021 |
| **Germany – NW** | Landesamt für Natur, Umwelt und Verbraucherschutz Nordrhein-Westfalen | | https://wolf.nrw/wolf/ | April 2021 |
| **Germany – SN** | Sächsisches Landesamt für Umwelt, Landwirtschaft und Geologie | | https://www.wolf.sachsen.de/ | April 2021 |
| **Germany – ST** | Landesamtes für Umweltschutz Sachsen-Anhalt | | https://lau.sachsen-anhalt.de/ | April 2021 |
| **Germany – SH** | Landesamt für Landwirtschaft, Umwelt und ländliche Räume | | https://www.schleswig-holstein.de/ | April 2021 |
| **Germany – MV** | Ministerium für Landwirtschaft und Umwelt | | https://wolf-mv.de/ | November 2021 |
| **Greece** | Hellenic Agricultural Insurance Organization | | Paraskevi Milioni | |
| **Italy** | Progetto LIFE WolfAlps | | Francesca Marucco* | |
| **Latvia** | Ministry of Agriculture | | Ilgvars Zihmanis | |
| **Lithuania** | State Service for Protected Areas under the Ministry of Environment | | Remigijus Uzdras | |
| **Netherlands** | Bij12 | | Angela van den Broek | |
| **Norway** | Naturvårdsverket | | https://rovbase.no/ | May 2021 |
| **Poland** | Regional Directorates for Environmental Protection | | Piotr Chmielewski* | |
| **Romania** | Ministry of Environment, Water and Forests | | Gavril Marius Berchi* | |
| **Slovakia** | Ministry of Environment | | Romana Uhrinová* | |
| **Slovenia** | Slovenia Forest Service | | Nives Pagon | |
| **Spain** | Regional Government of Asturias | | José Vicente López-Bao* | |
| **Sweden** | Naturvårdsverket | | Peter Jaxgard | |
| **Switzerland** | KORA Carnivore Ecology and Wildlife Management | | Fridolin Zimmermann | |

Table S.2: Livestock damage survey template.

| **Attribute** | **Field Type** | **Description** |
|---|---|---|
| CaseID | TEXT | Administrative File ID |
| Primary category | OPTION | Category of primary species affected, available levels: cattle, deer, dog, goat, horse, human, hunting dog, other, sheep, unspecified |
| Cause | OPTION | Concluding of cause of incident, available levels: disease, dog, fox, horned viper, mechanical injury, other, racoon, stillbirth, wolf |
| Assessment Level | OPTION | Level of assurance of assessment, available levels: assessment pending, no assessment possible, negative, uncertain, presumed correct, confirmed |
| Killed/Injured/Missing | NUMBER | Total number of animals killed, injured and missing |
| Num. Wolves | NUMBER | Number of wolves involved in the incident |
| Monetary Value | DECIMAL | Amount refunded to the claimant in EUR |
| Reported Date | DATE | Date of incident |
| Inspected Within Days | NUMBER | Num days between damage and inspection by authorities |
| Damage Prevention 1 | OPTION | Measure in place during incident, available levels: unspecified, unprotected (standard not met), closed electric fence, livestock guarding dog, euronet/wire fence, permanent shepherd, other qualified protection |
| Damage Prevention 2 | OPTION | as above |
| Damage Prevention 3 | OPTION | as above |
| Location | TEXT | Municipality address or geographical coordinates in WGS84 projection |
| Administrative Region | OPTION | Calculated based on location |
| Country | OPTION | Country |
| Data Constraints | TEXT | Comments on record quality |

Table S.3: Number of wolf-caused livestock damage incidents per year and per country (see main text). Countries marked with a * reported data only for a subset of the provinces with documented wolf presence. Countries marked with a + did not report an assessment level and we therefore analyzed all submitted records. For countries marked with a °, we analyzed compensated incidents.

| **Country** | **2018** | **2019** | **2020** |
|---|---|---|---|
| **Austria** | 34 | 20 | 61 |
| **Belgium** | 11 | 22 | 46 |
| **Croatia** | 1,111 | 1,229 | 1,541 |
| **Czech Republic**+ | 88 | 198 | 265 |
| **Estonia** | 121 | 178 | N/A |
| **Finland**° | 553 | 756 | 832 |
| **France**+ | 3,359 | 3,389 | 3,092 |
| **Germany** | 550 | 684 | 696 |
| **Greece** | 2,283 | 2,405 | 2,182 |
| **Italy***+ | 171 | 186 | 148 |
| **Latvia** | 30 | 39 | 22 |
| **Lithuania**+ | N/A | 408 | 492 |
| **Netherlands** | 44 | 25 | 95 |
| **Norway** | 703 | 239 | 205 |
| **Poland** | 680 | 727 | 121 |
| **Romania***° | 82 | 33 | 69 |
| **Slovakia**+ | 95 | 111 | 175 |
| **Slovenia** | 201 | 357 | 253 |
| **Spain***° | 2,145 | 2,474 | 2,237 |
| **Sweden** | 31 | 86 | 135 |
| **Switzerland** | 238 | 188 | 311 |
| **Total** | 12,530 | 13,754 | 12,978 |

Table S.4: Generalized linear models fitted to the average number of wolf-caused incidents excluding the outlier region of Asturias (ES120). The table shows the models with the best fit (delta AIC < 2.0) as well as the model including wolf presence only. The first three rows correspond to the Poisson model (P), for which all covariates except the factors were log-transformed. The lower six rows correspond to the Gaussian model (G), for which the average number of incident counts were power-transformed (see main text).

| | | Wolf presence | | Land cover classes | | | | | Factors | | | |
|---|---|---|---|---|---|---|---|---|---|---|---|---|
| Model | Inter. | *Per-A* | *Spo-A* | *P-A* | *NG-A* | *BLF-A* | *CF-A* | *MF-A* | *Prev* | *50ya* | *BIO11* | ΔAIC |
| P | -4.318e+01 | 1.997e-01 | 2.332e-02 | -1.675e-01 | 3.749e-01 | 1.274e-01 | 1.193e-01 | -8.794e-02 | + | + | 7.780e+00 | 0 |
| P | -4.076e+01 | 1.939e-01 | NA | -1.656e-01 | 3.699e-01 | 1.319e-01 | 1.302e-01 | -8.312e-02 | + | + | 7.366e+00 | 1.276e+01 |
| P | 9.963e-01 | 4.169e-01 | -3.073e-02 | NA | NA | NA | NA | NA | - | - | NA | 8.478e+03 |
| G | -6.854e+01 | 8.375e-04 | 1.000e-03 | -2.616e-03 | 1.963e-02 | NA | -1.190e-03 | NA | - | + | 2.572e-01 | 0 |
| G | -7.335e+01 | 7.741e-04 | 9.021e-04 | -2.563e-03 | 1.985e-02 | NA | -1.116e-03 | 5.891e-04 | - | + | 2.747e-01 | 3.369e-01 |
| G | -7.145e+01 | 6.094e-04 | 7.084e-04 | -2.946e-03 | 1.974e-02 | 1.234e-03 | -8.477e-04 | 8.339e-04 | - | + | 2.678e-01 | 7.851e-01 |
| G | -6.671e+01 | 7.724e-04 | 9.285e-04 | -2.807e-03 | 1.954e-02 | 5.805e-04 | -1.078e-03 | NA | - | + | 2.506e-01 | 1.665e+00 |
| G | -6.653e+01 | 8.175e-04 | 9.483e-04 | -2.529e-03 | 1.952e-02 | NA | -1.156e-03 | NA | + | + | 2.499e-01 | 2.501e+00 |
| G | 2.895e+00 | 8.106e-04 | 1.058e-04 | NA | NA | NA | NA | NA | - | - | | |

Table S.5: Generalized linear models fitted to the average number of wolf-caused incidents inculding the outlier region of Asturias (ES120). The table shows the models with the best fit (delta AIC < 2.0) as well as the model including wolf presence only. The first three rows correspond to the Poisson model (P), for which all covariates except the factors were log-transformed. The lower ten rows correspond to the Gaussian model (G), for which the average number of incident counts were power-transformed (see main text).

| | | Wolf presence | | Land cover classes | | | | | Factors | | | |
|---|---|---|---|---|---|---|---|---|---|---|---|---|
| Model | Inter. | *Per-A* | *Spo-A* | *P-A* | *NG-A* | *BLF-A* | *CF-A* | *MF-A* | *Prev* | *50ya* | *BIO11* | ΔAIC |
| P | -1.271e+02 | 2.048e-01 | -1.069e-01 | -8.594e-02 | 2.840e-01 | 1.999e-01 | 2.501e-01 | -1.867e-01 | + | + | 2.270e+01 | 0 |
| P | -1.366e+02 | 2.054e-01 | -1.044e-01 | NA | 2.668e-01 | 1.595e-01 | 2.271e-01 | -1.513e-01 | + | + | 2.438e+01 | 1.501e+02 |
| P | 9.480e-01 | 5.477e-01 | -1.735e-01 | NA | NA | NA | NA | NA | - | - | NA | 1.303e+04 |
| G | -3.070e+01 | 1.890e-04 | 2.102e-04 | NA | 6.114e-03 | 1.199e-03 | -2.520e-04 | 2.858e-04 | + | + | 1.174e-01 | 0 |
| G | -2.908e+01 | 2.451e-04 | 2.838e-04 | NA | 6.043e-03 | 9.888e-04 | -3.319e-04 | NA | + | + | 1.115e-01 | 3.979e-02 |
| G | -3.048e+01 | 7.937e-05 | 6.333e-05 | NA | 6.255e-03 | 1.573e-03 | NA | 4.056e-04 | + | + | 1.167e-01 | 7.292e-01 |
| G | -3.358e+01 | 2.052e-04 | 2.504e-04 | NA | 6.209e-03 | 1.149e-03 | -2.828e-04 | 3.087e-04 | - | + | 1.278e-01 | 1.023e+00 |
| G | -2.699e+01 | 7.006e-05 | NA | NA | 6.175e-03 | 1.700e-03 | NA | 5.542e-04 | + | + | 1.041e-01 | 1.231e+00 |
| G | -3.166e+01 | 2.662e-04 | 3.310e-04 | NA | 6.134e-03 | 9.245e-04 | -3.693e-04 | NA | - | + | 1.208e-01 | 1.436e+00 |
| G | -2.953e+01 | 1.948e-04 | 2.067e-04 | -3.459e-04 | 6.126e-03 | 1.264e-03 | -2.625e-04 | 2.946e-04 | + | + | 1.132e-01 | 1.649e+00 |
| G | -2.803e+01 | 2.516e-04 | 2.828e-04 | -2.965e-04 | 6.052e-03 | 1.039e-03 | -3.430e-04 | NA | + | + | 1.077e-01 | 1.806e+00 |
| G | -2.904e+01 | NA | NA | NA | 6.354e-03 | 1.866e-03 | 8.269e-05 | 5.410e-04 | + | + | 1.115e-01 | 2.008e+00 |
| G | 1.888e+00 | 3.669e-04 | 2.072e-05 | NA | NA | NA | NA | NA | - | - | NA | 2.483e+02 |
